# Movement shapes the structure of fish communities along a cross-shore section in the California Current

**DOI:** 10.1101/2021.09.16.458629

**Authors:** Jérôme Guiet, Daniele Bianchi, Olivier Maury, Nicolas Barrier, Fayçal Kessouri

## Abstract

Pelagic fish communities are shaped by bottom-up and top-down processes, transport by currents, and active swimming. However, the interaction of these processes remains poorly understood. Here, we use a regional implementation of the APex ECOSystem Model (APECOSM), a mechanistic model of the pelagic food web, to investigate these processes in the California Current, a highly productive upwelling system characterized by vigorous mesoscale circulation. The model is coupled with an eddy-resolving representation of ocean currents and lower trophic levels, and is tuned to reproduce observed fish biomass from fisheries independent trawls. Several emergent properties of the model compare realistically with observations. First, the epipelagic community accounts for one order of magnitude less biomass than the vertically migratory community, and is composed by smaller species. Second, the abundance of small fish decreases from the coast to the open ocean, while the abundance of large fish remains relatively uniform. This in turn leads to flattening of biomass size-spectra away from the coast for both communities. Third, the model reproduces a cross-shore succession of small to large sizes moving offshore, consistent with observations of species occurrence. These cross-shore variations emerge in the model from a combination of: (1) passive offshore advection by the mean current, (2) active swimming towards coastal productive regions to counterbalance this transport, and (3) mesoscale heterogeneity that reduces the ability of organisms to return to coastal waters. Our results highlight the importance of passive and active movement in structuring the pelagic food web, and suggest that a correct representation of these processes is needed for realistic simulations with marine ecosystem models.

## 1. Introduction

Marine ecosystems provide a variety of ecological services to humans, including food provision. These ecosystems are under increasing human pressure (Halpern et al. (2008), Duarte (2014)), which is degrading their state and could limit their ability to sustain fisheries (Blanchard et al. (2017)). Numerical models are essential tools to anticipate the evolution of marine ecosystems, integrating physical (Fox-Kemper et al. (2019)), biogeochemical (Bopp et al. (2013), Séférian et al. (2020)), and ecosystem components (Lotze et al. (2019)). Ecosystem models are particularly useful to parse key processes that structure the marine biosphere, and to explore the potential future states of the ocean under various emission (Blanchard et al. (2017)) and socio-economic scenarios (Maury (2017), Scherrer and Galbraith (2020)).

The dynamics of marine ecosystems reflects the interplay of physical, biological, and ecological processes, as well as direct human impacts through fishing. Understanding these processes is key to adequately represent and project the state of fisheries. Amongst ecological processes, size-dependent trophic interactions connect prey to predators (Estes et al. (2016)) and are modulated by the environment. Factors such as temperature and food abundance affect metabolism, growth, and reproduction, in turn altering the accumulation of biomass in the food web (Free et al. (2019)). Finally, oceanic currents and active swimming redistribute organisms and biomass, with effects that vary by species and size (Allen et al. (2018)). While the influence of food and temperature are generally accounted for in marine ecosystem models (Heneghan et al. (2021)), ocean currents and movement have received far less attention (Watson et al. (2015)).

Currents advect prey away from regions where primary production occurs (Popova et al. (2013), Messié and Chavez (2017)), and interactions between eddies and active swimming can enhance dispersion of organisms (Lévy et al. (2018)) but also lead to aggregations of mobile predators (Potier et al. (2014)), potentially increasing energy transfer to higher trophic levels (Woodson and Litvin (2015)). In coastal upwelling systems, ocean currents transport lower trophic levels away from the coast, driving a succession of pelagic communities from phytoplankton-dominated nearshore to zooplankton-dominated offshore (Keister et al. (2009), Messié and Chavez (2017)). The ability of mobile organisms to swim against currents can further modulate this transport (Drake et al. (2018)).

Here, we investigate how currents, including mesoscale eddies, interact with ecological processes to shape the distribution of higher trophic levels in the California Current, a highly dynamic eastern boundary upwelling system. The physical, biogeochemical and ecological processes occurring in the California Current have been largely studied (McClatchie (2014), Koslow and Davison (2016)). A considerable amount of data have been collected in this region, in particular with the CalCOFI program, which provides a wealth of in-situ observations from hydrography to biology (McClatchie (2014)). Trawl surveys of coastal pelagic species provide fisheries-independent observations of mid-trophic levels for the recent decades (Zwolinski et al. (2014)). Programs such as the Long Term Ecological Research Program and other scientific cruises add to this wealth of data, for instance sampling deep mesopelagic layers (Davison et al. (2013)). These observations facilitate the implementation and the validation of physical-biogeochemical models that reproduce ocean currents and primary production in the region, down to resolutions of kilometers or less (Gruber et al. (2006), Capet et al. (2008), Fiechter et al. (2018), Kessouri et al. (2020, 2021)). Models of species distribution, such as anchovy or sardines (Rose et al. (2015), Politikos et al. (2018)), and food web models (Horne et al. (2010), Kaplan et al. (2012), Koehn et al. (2016)) have also been implemented in this region. However, these models only include a limited representation of the interaction between oceanic currents and fish distribution.

Size-based marine ecosystem models allow a natural representation of both passive transport by currents and active swimming (Watson et al. (2015), Faugeras and Maury (2007)). They represent food web dynamics assuming that individual size is the fundamental structuring variable (Andersen and Beyer (2006), Maury and Poggiale (2013), Blanchard et al. (2017)). Size controls the propagation of biomass through the food-web, as indicated by the strong relationships between individual size and new biomass production (Brown et al. (2004)), reproduction (Kooijman (2010)), and predator-prey interactions (Barnes et al. (2010), Reum et al. (2019)). It also relates to swimming speed, both across species, and for different life stages within a species (Faugeras and Maury (2007)). Thus, size-structured models embedded within a realistic representation of ocean currents are ideal tools to test the relative influence of dynamical and environmental drivers on fish biomass distribution (Lefort et al. (2015), Le Mézo et al. (2016), Watson et al. (2015)).

We implement a regional configuration of such a size-based ecosystem model, The Apex Predators ECOSytem Model for the California Current (APECOSM-CC), and use it to investigate the interaction between movement and food web processes in the region. We discuss model formulation and parameterization in Section 2, together with observations from fishery-independent surveys used to constrain the model, and the procedure adopted to calibrate uncertain parameters. In Section 3 we present the model results, emphasizing the role of transport, and highlighting emergent model properties that we compare to observations. In Section 4 we discuss the results, focusing on the importance of ocean dynamics in establishing the cross-shore gradients in biomass and organism size that are observed in the California Current.

## 2. Materials and Methods

### 2.1. APECOSM in the California Current

#### 2.1.1. Overview of APECOSM

APECOSM links individual bioenergetic to the three-dimensional (3D) size-structured dynamics of biomass at the species and community levels, based on size-dependent elementary processes (Maury (2010), Maury and Poggiale (2013)). Here, it is used to simulate the biomass distribution of higher trophic levels, from hatching eggs (*L_min_* = 1 mm) to large predators (*L_max_* = 2 m), for two open-ocean communities: epipelagic and vertically migrating fish (see Fig. 1).

**Figure 1:**
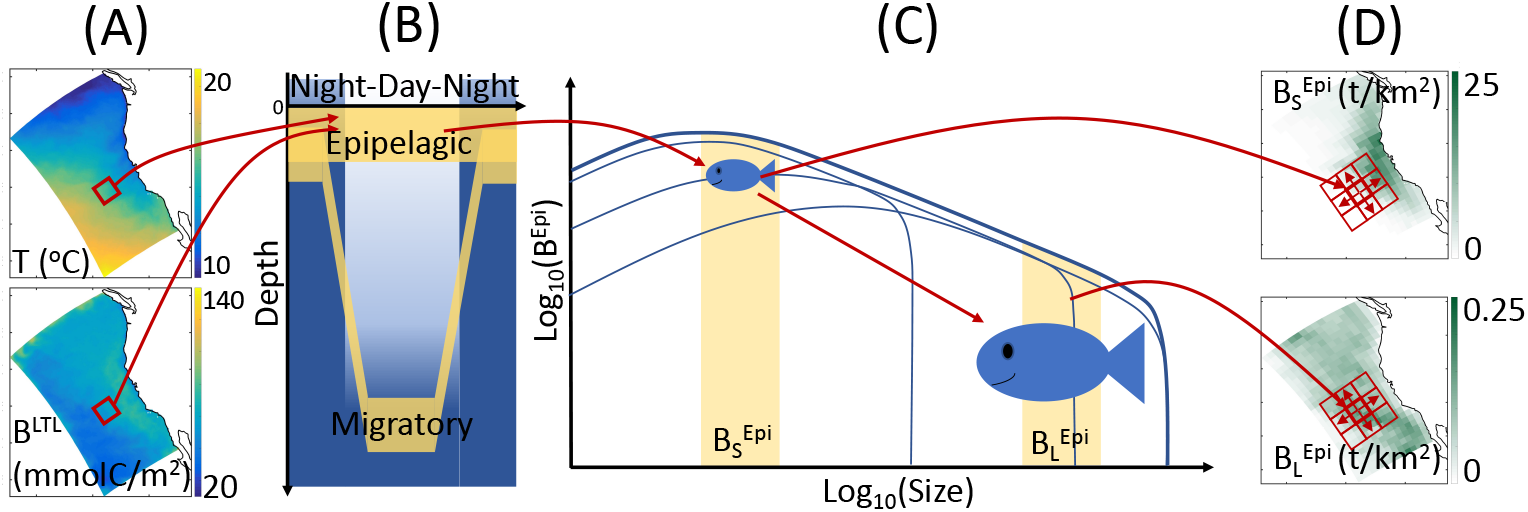
Schematic of APECOSM. (A) Illustration of the 3D environmental forcing from simulations with the physical-biogeochemical model ROMS-BEC, including temperature *T* and biomass of lower trophic levels (*B^LTL^*). (B) The 3D environmental forcing defines the habitat of distinct epipelagic and migratory communities. (C) Environmental conditions influence individual metabolism at different sizes, modulating growth, feeding, reproduction and mortality, thus driving the temporal evolution of the biomass density size-spectrum, here shown for epipelagic fish (*B^Epi^*). (D) The integrated biomass of different size ranges determines the spatial biomass distribution, here shown for small 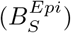 and large 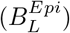 fish. Neighboring cells (red squares) exchange biomass by passive transport and active swimming.

Temperature (*T*), biomass of lower trophic levels (*B^LTL^*), dissolved oxygen concentration (*O*_2_), and photosynthetically available radiation (*PAR*) determine the daytime and nighttime preferred habitats of communities along the water column (see Fig. 1A,B). The temperature and food concentration determine the metabolism of individual fish (Fig. 1C). Size-dependent growth, maturation and reproduction are represented according to the Dynamic Energy Budget theory (DEB) (Kooijman (2010)). Small individuals feed on lower trophic levels, *B^LTL^*, while larger individuals feed on smaller prey. All rates vary with temperature following the Arrhenius equation. By integrating individual-level responses for species of different asymptotic size (*Lm*), APECOSM calculates the biomass density distribution of each species, i.e., the biomass per unit volume per unit size (*L*). Summing biomass density distributions for all species with *Lm* between 1 mm and 2 m (here discretized by 6 species with *Lm* = 0.1, 0.3, 0.6, 0.9, 1.4 and 2m), the model simulates the community size-spectrum (see Fig. 1C showing the biomass density distribution 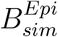 for the epipelagic group as example). Since larger individuals feed on smaller ones, different communities interact via predation when they co-occur in the water column. As shown in Fig. 1B, epipelagic and migratory fish co-occur during the night at the surface. Accordingly, fish feed within their communities separately during the day, and on both communities during the night.

Integrating the community size-spectra enables the calculation of the total biomass for different size ranges. Here, for diagnostic purposes, we subdivide the size spectrum into small (S, 0.03 < *L* < 0.3m), medium (M, 0.3 < *L* < 0.9m) and large (L, 0.9 < *L* < 2m) size classes. In the following, the biomass density of small epipelagic fish will be indicated by 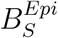, of medium epipelagic fish by 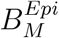, etc. (see Fig. 1C-D). Environmental properties and feeding interactions ultimately determine the biomass density distributions at each grid point (Fig. 1D).

The model accounts for movement by explicitly resolving passive advection by currents (with velocity 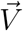), and active swimming between neighboring grid cells (Fig. 1D, the red squares). Swimming is modeled as a size-dependent advection-diffusion process that represents the migration of fish towards more favorable habitats (Faugeras and Maury (2007)).

#### 2.1.2. Physical and biogeochemical forcing

We force APECOSM-CC with a realistic three-dimensional (3D) ocean model simulation of the California Current based on the Regional Oceanic Model System (ROMS, Shchepetkin and McWilliams (2005)) coupled online with the Biogeochemical Elemental Cycling model (BEC, Moore et al. (2004)). This ROMS-BEC simulation has been run for the period 1997-2007 at a horizontal resolution of 4km (i.e., mesoscale eddy-resolving) that reproduces accurately the main patterns of physical and biogeochemical variability in the region (Renault et al. (2021), Deutsch et al. (2021)). Daily ROMS-BEC outputs on a coarsened grid (resolution of 16km) are used to drive APECOSM-CC. Physical forcings consist of temperature (*T*) and current velocities 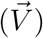. Biogeochemical forcings consist of dissolved oxygen (*O*_2_), *PAR* and the biomass of lower trophic levels *B^LTL^* that includes diatoms (*B^Diat^*), zoo-plankton (*B^Zoo^*), and particulate organic carbon (*B^POC^*) computed from particle flux *F_POC_* divided by a representative sinking velocity of 20*m/d*.

#### 2.1.3. Parameters and uncertainty

APECOSM-CC models pelagic fish in the California Current and relies on a total of 48 parameters and constants (see Table 1 and Supplementary Information S1). Many of these parameters are well constrained by the literature, while other remain to be specified. Among them, 5 parameters describe predator-prey interactions, and 6 control coupling with ROM-BEC (see Tab. 1). We determine them by testing the sensitivity of simulations on predetermined parameter ranges and choosing the values that produce the best match to observations (Section 3.1.2).

**Table 1:**
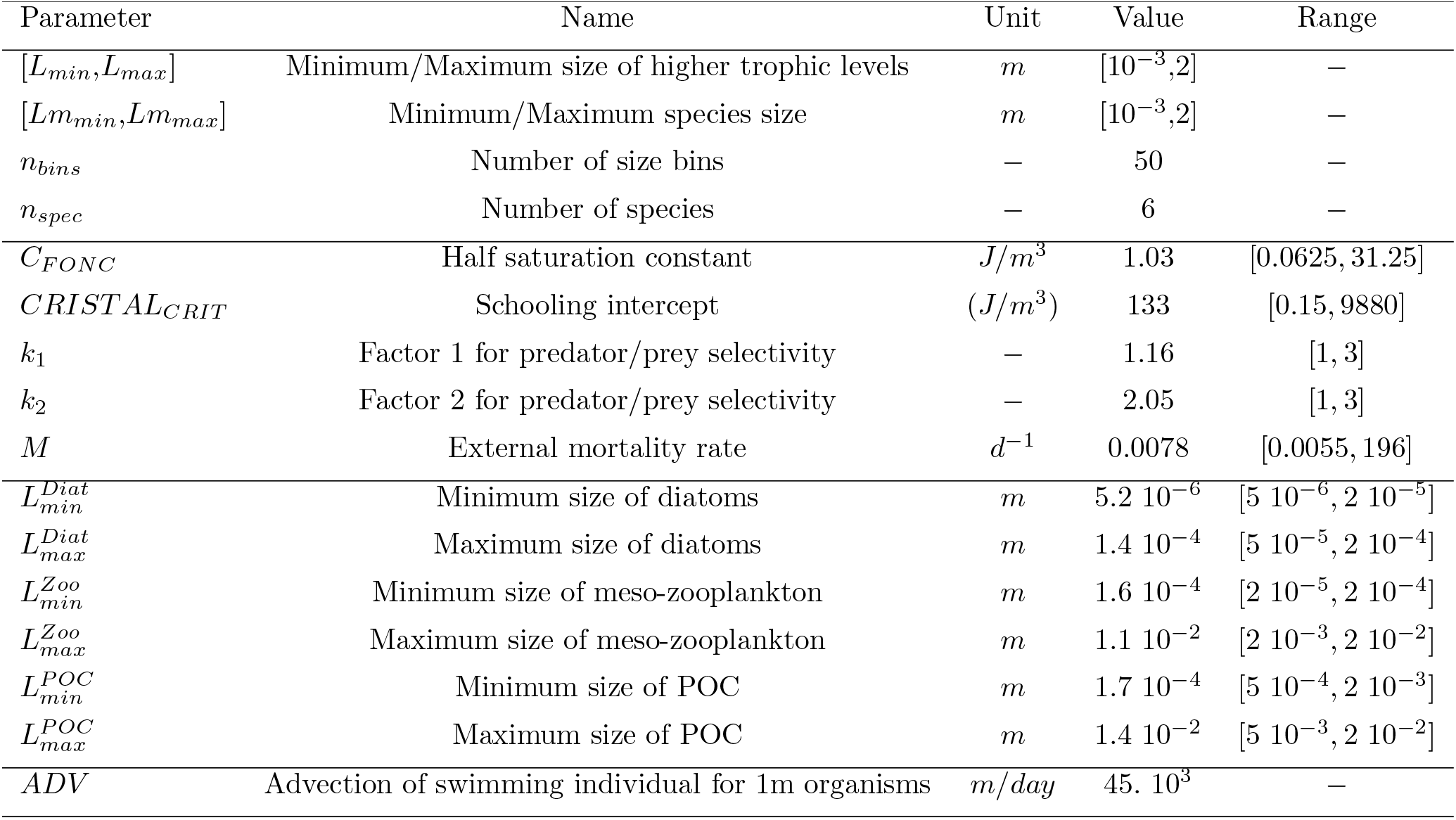
Parameters in APECOSM-CC referenced in the main text. See Supplementary Information S1 for full list.

| Parameter | Name | Unit | Value | Range |
| --- | --- | --- | --- | --- |
| $[L_{min}, L_{max}]$ | Minimum/Maximum size of higher trophic levels | $m$ | $[10^{-3}, 2]$ | — |
| $[Lm_{min}, Lm_{max}]$ | Minimum/Maximum species size | $m$ | $[10^{-3}, 2]$ | — |
| $n_{bins}$ | Number of size bins | — | 50 | — |
| $n_{spec}$ | Number of species | — | 6 | — |
| $C_{FONC}$ | Half saturation constant | $J/m^3$ | 1.03 | $[0.0625, 31.25]$ |
| $CRISTAL_{CRIT}$ | Schooling intercept | $(J/m^3)$ | 133 | $[0.15, 9880]$ |
| $k_1$ | Factor 1 for predator/prey selectivity | — | 1.16 | $[1, 3]$ |
| $k_2$ | Factor 2 for predator/prey selectivity | — | 2.05 | $[1, 3]$ |
| $M$ | External mortality rate | $d^{-1}$ | 0.0078 | $[0.0055, 196]$ |
| $L_{min}^{Diat}$ | Minimum size of diatoms | $m$ | $5.2 \cdot 10^{-6}$ | $[5 \cdot 10^{-6}, 2 \cdot 10^{-5}]$ |
| $L_{max}^{Diat}$ | Maximum size of diatoms | $m$ | $1.4 \cdot 10^{-4}$ | $[5 \cdot 10^{-5}, 2 \cdot 10^{-4}]$ |
| $L_{min}^{Zoo}$ | Minimum size of meso-zooplankton | $m$ | $1.6 \cdot 10^{-4}$ | $[2 \cdot 10^{-5}, 2 \cdot 10^{-4}]$ |
| $L_{max}^{Zoo}$ | Maximum size of meso-zooplankton | $m$ | $1.1 \cdot 10^{-2}$ | $[2 \cdot 10^{-3}, 2 \cdot 10^{-2}]$ |
| $L_{min}^{POC}$ | Minimum size of POC | $m$ | $1.7 \cdot 10^{-4}$ | $[5 \cdot 10^{-4}, 2 \cdot 10^{-3}]$ |
| $L_{max}^{POC}$ | Maximum size of POC | $m$ | $1.4 \cdot 10^{-2}$ | $[5 \cdot 10^{-3}, 2 \cdot 10^{-2}]$ |
| $ADV$ | Advection of swimming individual for 1m organisms | $m/day$ | $45 \cdot 10^3$ | — |

First, predator-prey interactions depend on the rates of prey encounter and handling by predators. The half saturation constant (*C_FONC_*) and schooling threshold (*CRISTAL_CRIT_*) set the intensity of these interactions. The half saturation constant represents the limitation of biomass ingestion by encounter or assimilation rates, and falls within the range [0.0625, 31.25] *J/m*^3^ (see Supplementary Information S2). The schooling threshold determines the relative biomass density above which 50% of prey become accessible to predation, the remaining 50% assumed to be too dispersed to be effectively preyed upon (Maury (2017)). Based on the minimum and maximum biomass densities of lower trophic levels (*B^LTL^*) we select it within the range [0.15, 9880] *J/m*^3^ (see Supplementary Information S3).

Second, feeding follows a selectivity function that depends on the predatorprey mass ratio (*PPMR*) and selectivity width (*σ*). In APECOSM-CC, both parameters are controlled by two constants, *k*_1_ and *k*_2_, that we set to be within the range [1, 3], leading to a predator-prey mass ratio within [245, 19260] for a 0.25m long predator, and a selectivity width within [1.0, 1.7] (see Supplementary Information S4 for details).

Third, the model accounts for predation mortality, ageing, and starvation. A density-dependent background mortality is included to account for external sources of mortality (*M*), including disease and predation by functional groups not explicitly represented by the model. This additional mortality is selected to produce a mortality timescale for the smallest individuals of a population of between 1 week and 6 months, resulting in a mortality within the range of [0.0055, 196] *d*^−1^ (see Supplementary Information S5).

Finally, the size range of lower tropic levels influences the coupling of APECOSM-CC with ROMS-BEC by setting the intercept of the prey size spectrum feeding small predators (see Supplementary Information S6). It is controlled by 6 configuration parameters that determine the size of smallest and largest diatom cells, zooplankton organisms, and POC 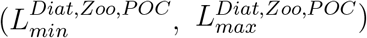. We allow slight variations of these minimum and maximum sizes around default values (see Tab. 1).

### 2.2. Pelagic fish in the California Current

#### 2.2.1. Surface- and Mid-water trawls

To tune and evaluate APECOSM-CC, we collected data from approximately 1700 fisheries independent trawls from long-running research and monitoring programs in the California Current (Davison et al. (2013), McClatchie (2014), Zwolinski et al. (2014)), extending from the epipelagic to the mesopelagic zones.

The Surface-Water (SW) coastal pelagic species surveys (Zwolinski et al. (2014)) provide 1556 distinct trawl-based biomass estimates from March to October between 2003 and 2016 (Fig. 2A). Most of these surveys consist of nighttime surface trawls, targeting fish aggregations with Nordic 264 trawls, covering most of the California Current, with higher occurrence nearshore (Fig. 2B). We use the surface-water trawls to estimate the observed biomass density distribution of epipelagic species 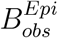 in the size range [0.03, 2.]m. Surface-water trawls also provide species composition for each trawl. Combining the weight per species sampled (*w_s_*) and the species asymptotic sizes (*Lm_s_*) from Fishbase (Froese and Pauly (2016)), we estimate the mean asymptotic fish length for each trawl 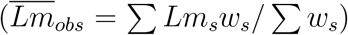. We compute 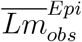 for pelagic species with a depth range shallower than 150m (based on Fishbase); 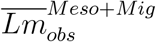 for vertically migrating species, i.e., migratory pelagic fish with a depth range deeper than 150m, and mesopelagic fish (see Section 3.2.1). Finally, for a fraction of the surface-water trawls the length of each individual (*L*) is also measured. We use these to determine the abundance of individuals of different size in logarithmically equal bins and compare this distribution, the size spectrum, with simulations (see Section 3.2.2 and Supplementary Information S7 for more details).

**Figure 2:**
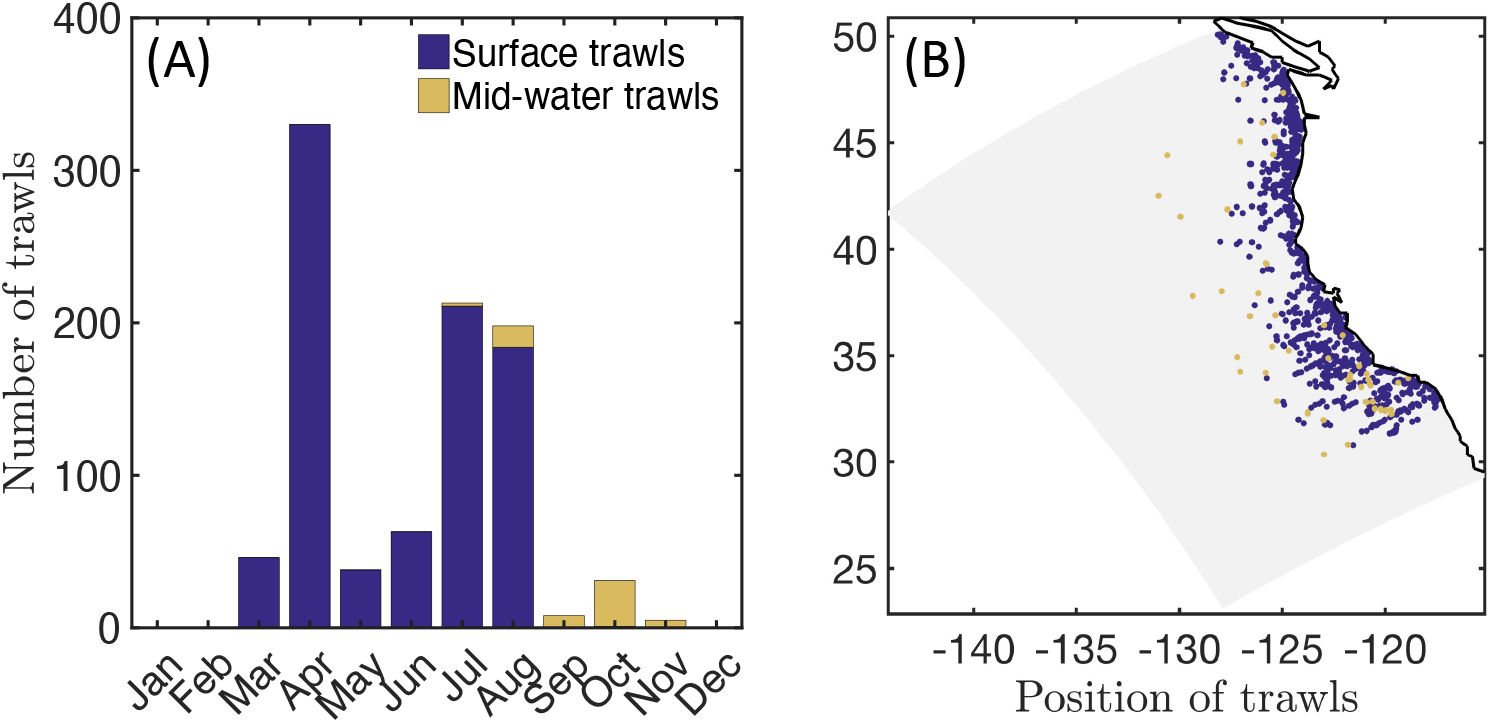
Trawl data in the California Current. (A) Total number of fisheries independent trawls per month. (B) Spatial distribution of fisheries independent trawls (blue, surface trawls; orange, mid-water trawls) and focus region for fish biomass time-series reconstruction (red).

In addition, Mid-Water (MW) trawls from the Seaplex, Orca-Wale and CCE cruises from August to November in 2008 and 2009 (Fig. 2A), are used to estimate the biomass density distribution in deep waters (Davison et al. (2013)). These trawls are conducted during the day and at night, down to different depths, with Isaacs-Kid trawls (IKMT, Orca-Wale, 95 trawls) or Matsuda-Oozeki-Hu trawls (MOHT, Seaplex and CCE, 55 trawls) and target centimeter-size forage and mesopelagic fish. The trawls cover the entire California Current, from coastal to offshore waters (Fig. 2B). We use the mid-water trawls to estimate the observed biomass density distribution of vertically migrating species 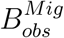 in the size range [0.0017, 0.1]m (see Davison et al. (2013) for the biomass estimates).

To complement the trawl observations, we use the biomass reconstruction by Koslow and Davison (2016) of the spawning stock biomass for epipelagic and mesopelagic planktivores from 1951 to 2011 in the Southern California Current (see region in Fig. 2B). We use these reconstructions to obtain estimates of the average, minimum, and maximum ratio of epipelagic to migratory mesopelagic biomass, resulting in 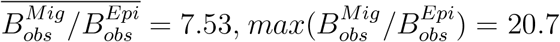, and 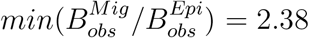 respectively.

#### 2.2.2. Species occurrence and asymptotic lengths

Occurrence data obtained from the Ocean Biodiversity Information System database (OBIS) are used to inform the spatial distribution of species of increasing size in the California Current. We compiled spatial occurrence along with life-stage and species identity for surface epipelagic fish, vertically migrating predators, and mesopelagic fish respectively accounting for 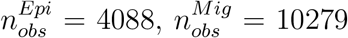 and 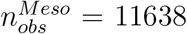 occurrences each. From the species identity, we associated each occurrence to the species asymptotic size (*Lm*, using Fishbase, Froese and Pauly (2016)) and determined the proportion of small (0.04 < *Lm* < 0.4m), medium (0.4 < *Lm* < 0.9m) and large (0.9 < *Lm* < 2m) species as a function of the distance d from the coast, calculated along regular bins of width *dx* = 33km. This cross-shelf probability of occurrence of species of increasing asymptotic size provides an observational estimate of the distribution of different sizes classes in the California Current (see Section 3.3.1 and Supplementary Information S8).

### 2.3. Simulation and analysis

#### 2.3.1. Implementation and uncertainty

To constrain the 11 undetermined model parameters, we develop a calibration procedure on a simplified one-dimensional (1D) configuration of the domain, here referred to as APECOSM-1D, that reduces spatial variability to 15 independent 1D stations representative of 15 eco-regions in the California Current (see Fig. 3A and Supplementary Information S9). At each station, we drive APECOSM-1D with environmental forcings based on simulated year 2001 from ROMS-BEC, averaged over the eco-region (Fig. 3B). Each station is spun-up for 100 years to reach a stable seasonal cycle, and is then analyzed. The model timestep for predatory interactions, growth, and mortality is one day, averaging daytime and nighttime conditions. This simplified configuration disregards the role of horizontal movement, but allows narrowing down parameter combinations that produce realistic solutions. We run 5000 replicates of this APECOSM-1D configuration applying a Monte Carlo approach, each with a distinct combinations of the 11 undetermined parameters randomly chosen from prior distributions (see Fig. 3C, and Table 1).

**Figure 3:**
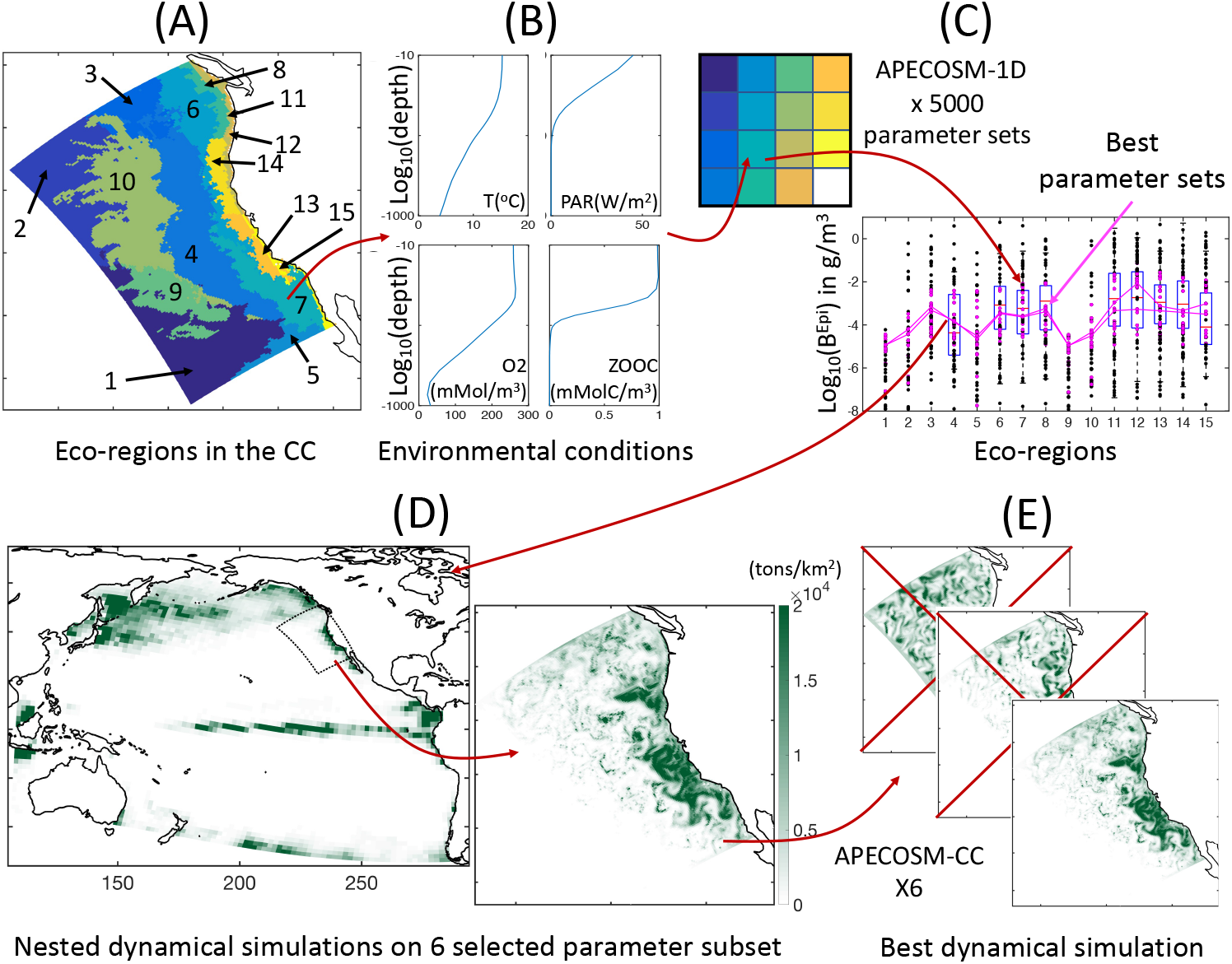
Schematic of the optimization approach. (A) Eco-regions of the California Current used as independent stations for model tuning. (B) Illustration of averaged vertical profiles used to force the model in each station, here station #7. (C) Illustration of the Monte Carlo procedure, whereby each eco-region is reduced to a single grid cell in APECOSM-1D, 5000 replicates of this 1D configuration are run, and optimal parameter sets are selected by comparison with observations. (D) Illustration of the parent Pacific and nested California Current 3D dynamical simulations. (E) Selection of the best regional 3D dynamical simulation from 6 optimal simulations with different parameter sets.

By comparing the ensemble of APECOSM-1D simulations to observations, we identified 6 best plausible parameter sets (see Section 3.1.1). With these we run 6 replicates of the 3D APECOSM-CC model forced with coarsened 16km resolution output from ROMS-BEC. Biomass transport inside and outside the domain is resolved by “nesting” the California Current configuration within a parent simulation for the Pacific Ocean run at 64km resolution, with forcing from a Pacific ROMS-BEC configuration (see Fig. 3D). We use the parent APECOSM simulation to provide boundary conditions for fish biomass to the California Current domain, adopting a two-dimensional boundary radiation scheme (Marchesiello et al. (2001)). These simulations are span-up with repeated 2001 annual forcing for 100 years and resolve spatial transport with a timestep of 20 minutes for numerical stability.

To analyze the role of transport in structuring marine ecosystems, we keep the 3D simulation that best reproduces biomass densities in eco-regions (see Fig. 3E), and then force it with two inter-annual cycles of 10 years (1997 to 2006). We analyze the last 8 years from weekly outputs. To account for the uncertainty introduced by undetermined model parameters in this simulation, we run 10 replicates with slight parameter variations, +/ − 10% for food web parameters, and ×3 or ×1/3 for movement parameters.

#### 2.3.2. Diagnostic features of fish communities

At each grid cell, APECOSM calculates the biomass density 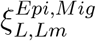 per size class *L*, per species asymptotic size *Lm*, for the epipelagic and migratory communities. For comparison with observation and analysis, we sum biomass over various combinations: 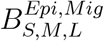 the biomass of small, medium and large fish; 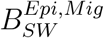 and 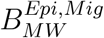 the biomass over size ranges respectively matching surface- and mid-water trawls selectivities; *B^Epi,Mig^* the total biomass density per community (see Table 2 for the equations). We also characterize the relative distribution of small, medium and large fish groups by analyzing biomass along a cross-shore section. This cross-shore distribution is estimated by averaging the biomass density 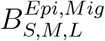 in regular spatial distance bins *i* (*dx_i_* = 33km) from the coast to the open ocean. Furthermore, for sensitivity analysis the “spread” of the biomass distribution in the region is summarized by the distance from coast 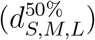 within which 50% of the total biomass in the region 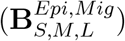 occurs (see Table 2).

**Table 2:** Metrics for the analysis of ecosystem features.

| Metric | Description | Definition | Unit |
| --- | --- | --- | --- |
| $B_{SW}^{Epi,Mig}$ | Biomass for surface water (SW) trawl selectivity | $\int_{0.03}^2 \tilde{\xi}_{L,Lm}^{Epi,Mig} dL$ | in $g/m^2$ |
| $B_{MW}^{Epi,Mig}$ | Biomass for mesopelagic water (MW) trawl selectivity | $\int_{0.0017}^{0.1} \tilde{\xi}_{L,Lm}^{Epi,Mig} dL$ | in $g/m^2$ |
| $B_S^{Epi,Mig}$ | Biomass of small size fish | $\int_{0.03}^{0.3} \tilde{\xi}_{L,Lm}^{Epi,Mig} dL$ | in $g/m^2$ |
| $B_M^{Epi,Mig}$ | Biomass of medium size fish | $\int_{0.3}^{0.9} \tilde{\xi}_{L,Lm}^{Epi,Mig} dL$ | in $g/m^2$ |
| $B_L^{Epi,Mig}$ | Biomass of large size fish | $\int_{0.9}^2 \tilde{\xi}_{L,Lm}^{Epi,Mig} dL$ | in $g/m^2$ |
| $B^{Epi,Mig}$ | Total fish biomass | $\int_{0.001}^2 \tilde{\xi}_{L,Lm}^{Epi,Mig} dL$ | in $g/m^2$ |
| $\mathbf{B}_{S,M,L}^{Epi,Mig}$ | Total fish biomass in the CC domain | $\int_{CC} B_{S,M,L}^{Epi,Mig} dS$ | in $g$ |
| $d_{S,M,L}^{50\%}$ | Distance from coast within which 50% of $\mathbf{B}_{S,M,L}^{Epi,Mig}$ occurs | - | in $km$ |
| $L_{cut}^{Epi,Mig}$ | Size of largest individuals per community | Size class $L_i$ where $\tilde{\xi}_{L_i}^{Epi,Mig} / \tilde{\xi}_{L_{i+1}}^{Epi,Mig} > 3$ | in $m$ |
| $\lambda^{Epi,Mig}$ | Slope of the community spectrum | Slope of the distribution $\tilde{\xi}_L^{Epi,Mig}$ over $[0.02, L_{cut}]$ | - |
| $\Delta_{META}^{Epi,Mig}$ | Source/Sink of biomass due to metabolism | $(G - M)_{S,M,L}^{Epi,Mig}$ | in $g/m^2/d$ |
| $\Delta_{CURR}^{Epi,Mig}$ | Source/Sink of biomass due to passive transport by currents | $\nabla F_{Phys\ S,M,L}^{Epi,Mig}$ | in $g/m^2/d$ |
| $\Delta_{SWIM}^{Epi,Mig}$ | Source/Sink of biomass due to active transport by swimming | $\nabla F_{Swim\ S,M,L}^{Epi,Mig}$ | in $g/m^2/d$ |

The size of the largest individuals sustained in each community is an emergent property of the model. We compute the maximum length reached in each numerical cell, 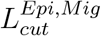, for the epipelagic and migratory communities, as the size at which the community-level biomass density distribution 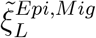 “breaks”, i.e., the size above which biomass rapidly drops (see Table 2). Because the community-level biomass size spectrum roughly follows a power law, we use the power law slope *λ^Epi,Mig^* between the minimum (*L* = 2 cm) and maximum size (*L_cut_*) as a metric of the relative abundance of small and large individuals.

Finally, the model allows analyzing the mechanisms that lead to local biomass accumulation or reduction and how these relate to spatial transport. We extract net sources and sinks of biomass in each grid cell, as driven by the following processes: local growth and mortality 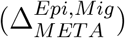; transport by currents 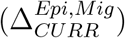; transport by swimming 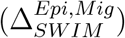 (see Table 2 and Supplementary Information S10 for the growth *G*, mortality *M*, and biomass fluxes *F_Phys,Swim_*). Their cross-shore distribution is estimated like for the biomass.

## 3. Results

### 3.1. Biomass distribution in the California Current

#### 3.1.1. Selection of a best simulation

The 5000 simulations with APECOSM-1D lead to a variation of several orders of magnitude for epipelagic 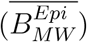 and migratory 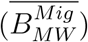 biomass, once averaged over the 15 eco-regions (see Fig. 4B). Only 16% reproduce the observed biomass density from mid-water trawls, 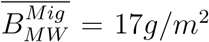, +/−1 order of magnitude (dotted horizontal lines Fig. 4B). The +/−1 order of magnitude range corresponds to the observed variation between the less productive offshore trawls and the trawls in the coastal upwelling (see observations Fig. 5). A smaller fraction (3%) also capture the observed ratio of mesopelagic to epipelagic biomass, namely 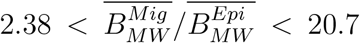 (oblique lines Fig. 4B, see Supplementary Information S11 for the sensitivity of APECOSM-1D).

**Figure 4:**
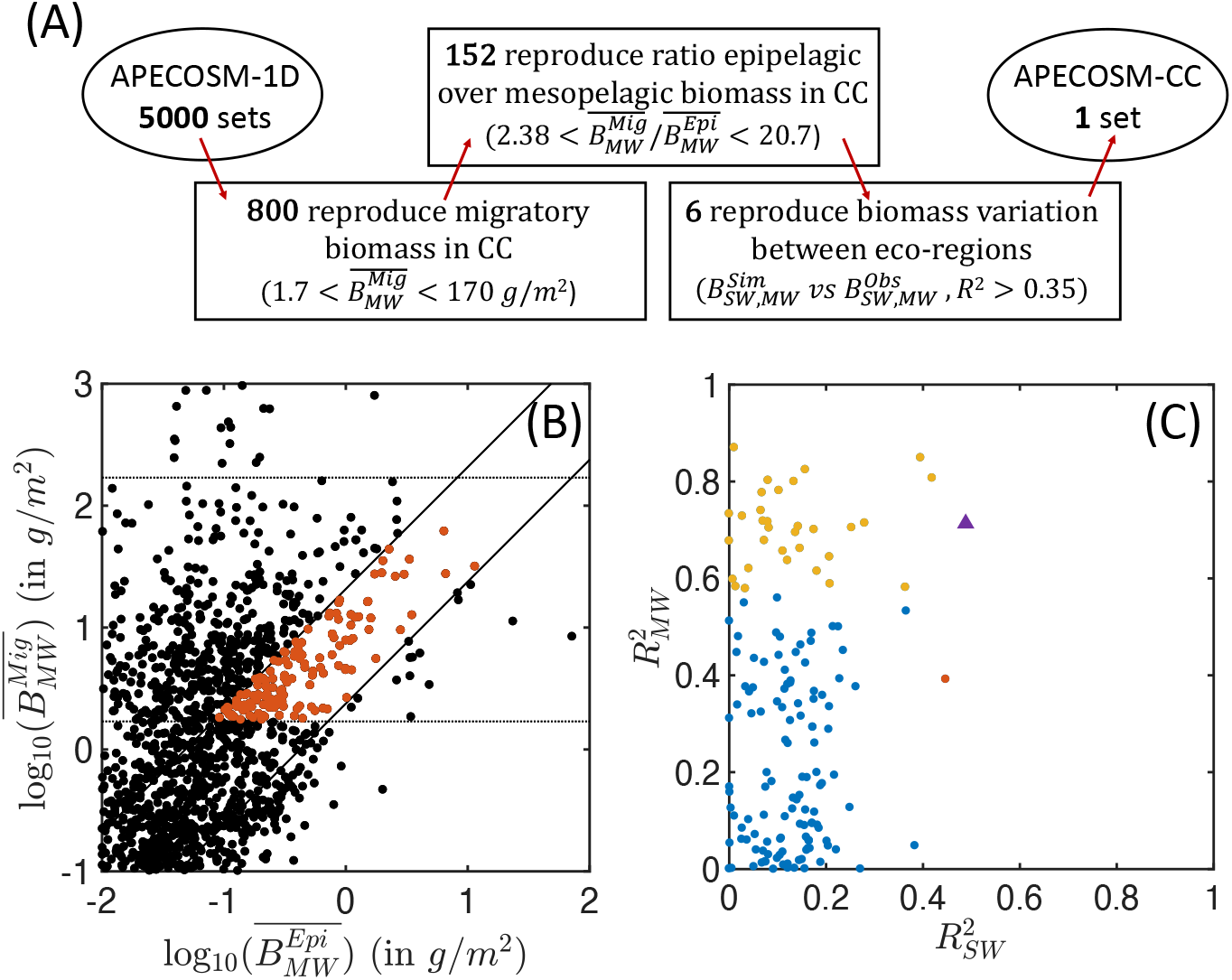
Summary of the ensemble of simulations with APECOSM-1D. (A) Schematic of the steps used to identify the optimal parameter sets. (B) Average migratory biomass density 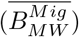 as a function of average epipelagic biomass density 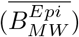. (C) Coefficient of determination of the correlation between observed and simulated 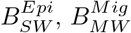, in the 15 eco-regions, for the best ensembles shown in red in panel (B). In panel (B) the dotted horizontal lines show the acceptable range for migratory biomass densities. The solid diagonal lines show the limits of acceptable migratory to epipelagic biomass ratio based on observations (between 2.38 and 20.7). In panel (C) the red and yellow dots corresponds to ensembles for which epipelagic and mesopelagic biomass respectively correlate significantly with observations (*p* < 0.05) (note that there is only one red dot). Blue dots are ensembles for which there is no statistically significant correlation between simulations and observation. The purple triangle corresponds to the final best set of parameters selected for the 3D APECOSM-CC dynamic simulations.

**Figure 5:**
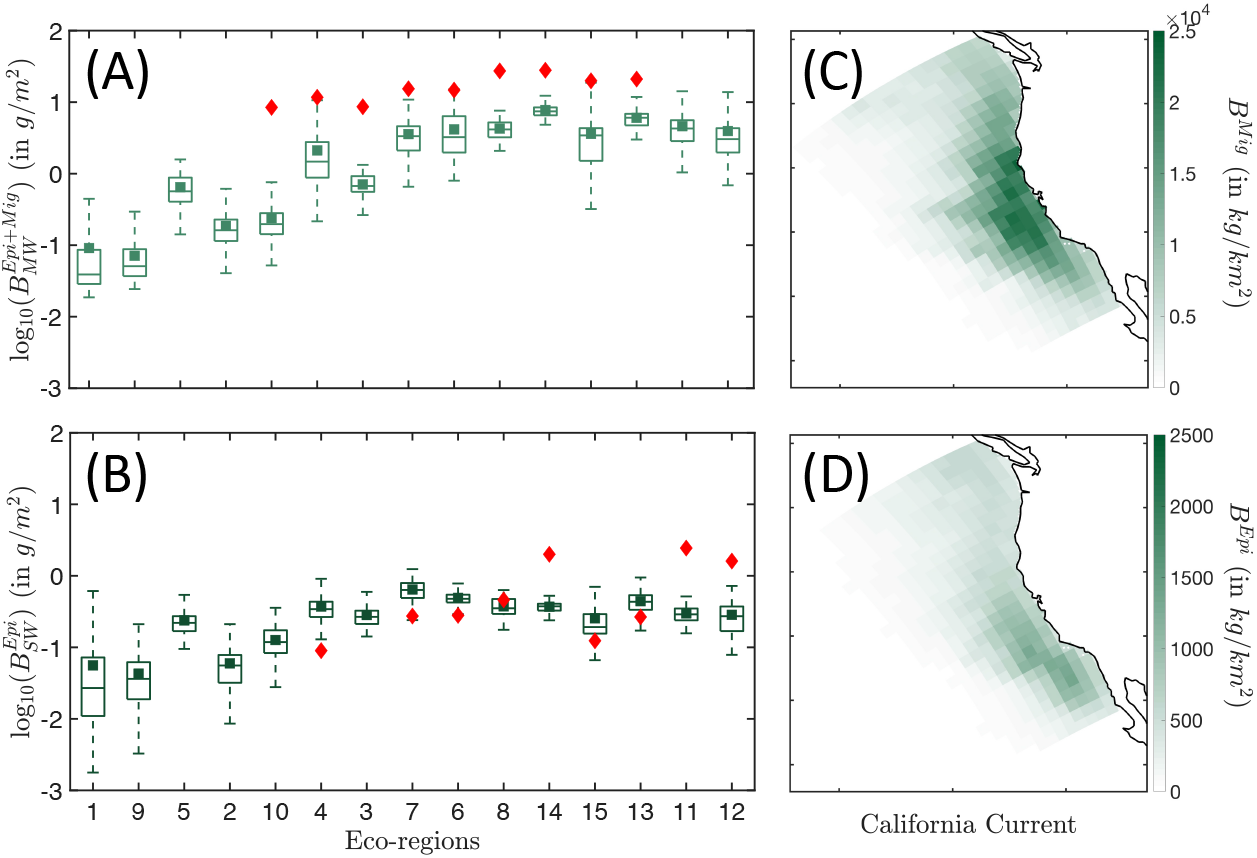
Biomass distribution in the California Current. (A) Simulated (light green) and observed (red) biomass densities *B^Epi+Mig^* per eco-regions for the mid water (MW) trawls. (B) Simulated (dark green) and observed (red) biomass densities *B^Epi^* per eco-regions for the surface water (SW) trawls. (C) Spatial distribution of the annual mean biomass density for the migratory community *B^Mig^* binned in Δ*x* = 96km grid cells. (D) Spatial distribution of the annual mean biomass density for the epipelagic community *B^Mig^*, binned in Δ*x* = 96km grid cells. The simulation results in panels (A-B) are determined from the average of years 1999 to 2006. Eco-regions are ranked from low to high primary production. In the box-plot the central mark indicates the median, the bottom and top edges of the box indicate the 25*^t^h* and 75*^t^h* percentiles respectively, while the whiskers extend to the most extreme data points that are not considered to be outliers. The square marker is the averaged biomass density per eco-region, and the diamond the averaged observed biomass density per eco-region.

Out of these 152 simulations (shown in red in Fig. 4B), an even smaller fraction captures the biomass variation between eco-regions. Figure 4C shows the Pearson correlation of annual mean simulated biomass densities compared to observations for the epipelagic community (compared with surface-water trawls) and the migratory community (compared with mid-water trawls). Analysis of these correlations shows that different parameter sets can properly capture variations in epipelagic (red markers in Fig. 4C, *p* < 0.05), migratory (yellow markers, *p* < 0.05), or both communities (purple marker) at statistically significant levels. We select 6 sets of parameters with higher coefficient of determination for both the epipelagic and migratory biomass densities 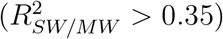 for fully dynamical simulations with APECOSM-CC (see Supplementary Information S11 for the parameters).

With transport, 3 out of 6 APECOSM-CC simulations maintain a realistic ratio between epipelagic and migratory biomass. We keep the simulation that best reproduces the regional biomass density gradients observed between ecoregions in the California Current (triangle marker in Fig. 4; see Table 1 for best parameters; see Supplementary Information S12 for 6 APECOSM-CC simulations).

#### 3.1.2. Observed vs. simulated biomass distribution

Observation show an increase in biomass density in both the surface (SW) and mid-water (MW) layers when moving from less productive offshore waters (Fig. 5A,B, red diamonds in eco-regions #3, 4, 6, 7, 10) to more productive coastal waters (#8, 11, 12, 13, 14, 15). Furthermore, seasonal eco-regions at higher latitudes (#6, 8, 11, 12, 14) hold comparatively higher biomass densities than more equatorward eco-regions (#7, 13, 15).

The spatial distribution of simulated biomass density corresponding to mid-water trawls 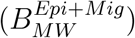, averaged from July to November, reproduces observed patterns (*R*^2^ = 0.76 for the correlation between model and observations, see Fig. 5A). However, the model underestimates observations (green squares Fig. 5A), with modeled biomass densities on average 29% lower than mid-water trawls. Spatially, the biomass density of vertically migrating fish *B^Mig^* is higher on the shelf and in the core of the upwelling, and much smaller offshore (Fig. 5C).

For surface-water trawls, when averaged from March to August, regional variations of simulated biomass densities for epipelagic fish are poorly captured (*R*^2^ = 0.07). This weak correlation mostly originates from a strong underestimate of the biomass density 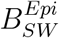 in the coastal regions (#11, 12, 14 Fig. 5B). Away from the coast, regional averages match more closely the surface-water trawls. Similar to migratory biomass, epipelagic biomass also accumulates along the coast, but is higher towards the southern limb of the upwelling and the Southern California Bight (Fig. 5D).

Overall, dynamic simulation leads to a coherent distribution of migratory biomass, and a more irregular distribution of pelagic biomass. In the southern California Current (red box in Fig. 2), the average ratio of pelagic to migratory biomass is well within the range of observations, i.e., 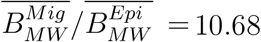.

#### 3.1.3. Sensitivity to parameters uncertainty

A parameter sensitivity analysis with the best simulation indicates that the predation parameters *k*_1,2_ exert the strongest controls on biomass accumulation 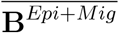 (Fig. 6). Larger predator-prey mass ratio (*k*_1_ – 10% or *k*_2_ + 10%) increase biomass accumulation in the domain. This accumulation benefits the migratory community, such that 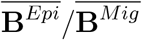 decreases. The latter also reaches smaller size 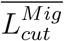. Thus, the biomass of larger individuals decreases to the benefit of smaller ones (see variations of 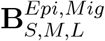). Opposite responses are observed for smaller predators-prey mass ratios (*k*_1_ + 10% or *k*_2_ – 10%). The cross-shore biomass distribution is slightly impacted by variations of the predation parameters (see changes in 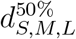).

**Figure 6:**
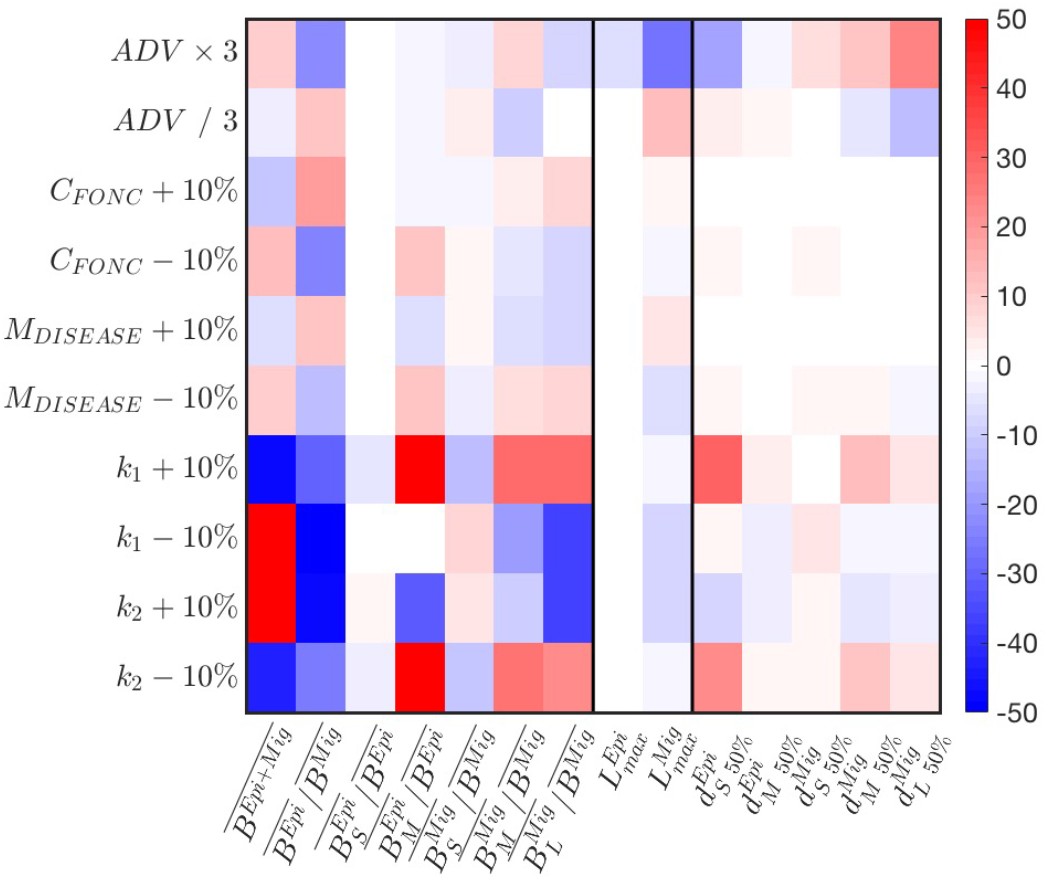
Sensitivity of APECOSM-CC to food-web parameters. We test variations around +/−10% for the food-web parameters, × / 3 for the parameters controlling movement. The sensitivity is expressed as % variation compared to the reference simulation with the best parameter set.

A decrease of prey searching rate, obtained by increasing the half saturation constant *C_FONC_*, and the increase of mortality *M*, both decrease biomass accumulation, while benefiting epipelagic fish, such that 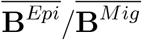 increases. While the effects translate to variations of the relative abundance of small, medium and large fish (see 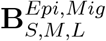), they have almost no impacts on the spatial biomass distribution (see 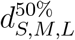). Opposite responses are observed for increasing searching rate *C_FONC_* and decreasing *M*.

Finally, a faster swimming speed (*ADV*) allows increasingly large migratory fish to expand their distribution offshore, while epipelagic fish tend to cluster nearshore (see changes in 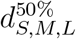 in Fig. 6). This drives an accumulation of biomass that benefits epipelagic species, such that 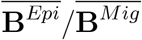 increases, while the maximum size of migratory species 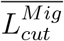 decreases.

Sensitivity tests shown in Fig. 6 reveal that most features of a fully dynamical simulation are similarly or less impacted than the amplitude of variation applied to a single parameter, suggesting non-linear, compensatory effects in the model. Except for the predator-prey selectivity parameters *k*_1,2_, most variations are within the +/ − 10% or 1/3 to 3 × range. Thus, fully dynamical simulations and the following analysis are robust to small parameter variations.

### 3.2. Emergent features of fish communities

#### 3.2.1. Community maximum sizes

The average fish size from trawl observations 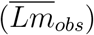 is different for epipelagic and migratory species (Fig. 7 and Supplementary Information S7). Epipelagic species are mostly smaller, with asymptotic sizes between 0.165m and 3.3m, and a mean of 0.4m (Fig. 7A in blue). Migratory species exhibit a larger range, from 0.077m to 4m, with two modes, the first around 0.2m when migratory mesopelagic fish dominate the trawls, and the second around 1m when large predators are captured (Fig. 7B in purple).

**Figure 7:**
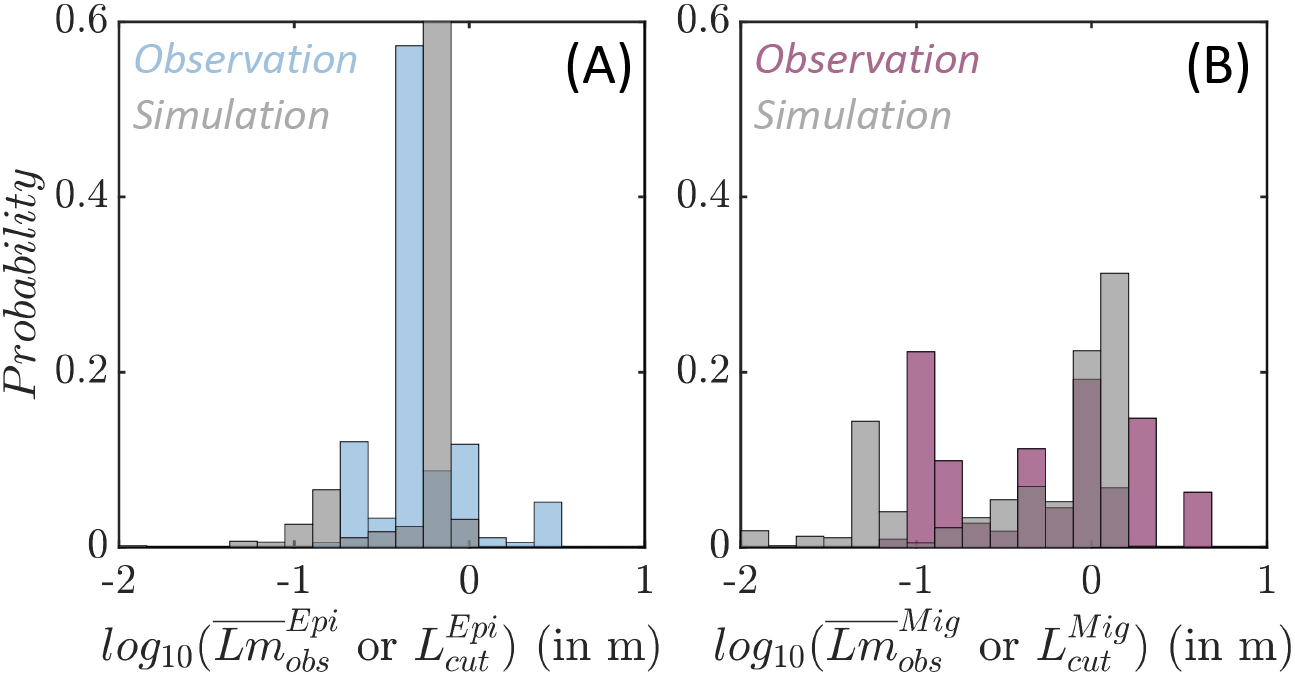
Species size distributions. (A) Observed average species size per trawl 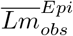 (blue) and simulated maximum size per grid cell (grey) for the epipelagic community 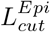. (B) Observed average species size per trawl 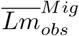 (purple) and simulated maximum size per grid cell (grey) for the migratory community 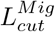.

In APECOSM-CC, the environment determines the maximum size of the species sustained in each grid-cell for each community. The maximum size per community 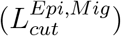 simulated in each grid cell of the domain matches observed size differences. Epipelagic species are smaller, with a mean maximum size of 0.62m (Fig. 7A in grey). Migratory species reach larger sizes, up to 1.3m at most locations (Fig. 7B in grey). The sensitivity analysis shown in Fig. 6 indicates that this maximum size is sensitive to the predator-prey selectivity range (*k*_1,2_), as well as parameters controlling swimming (*ADV*).

#### 3.2.2. Size-spectra slopes

In the simulations, the biomass density distribution closely follows a power law (see the average size spectra in Fig. 8A). Regional variations in the power law slope (λ) reveal variations of the biomass of large individuals relative to small ones. Note that here slopes are expressed as a function of fish weight or volume, derived from size following *V* = (*δL*)^3^, with *δ* a shape factor (see Supplementary Information S1).

**Figure 8:**
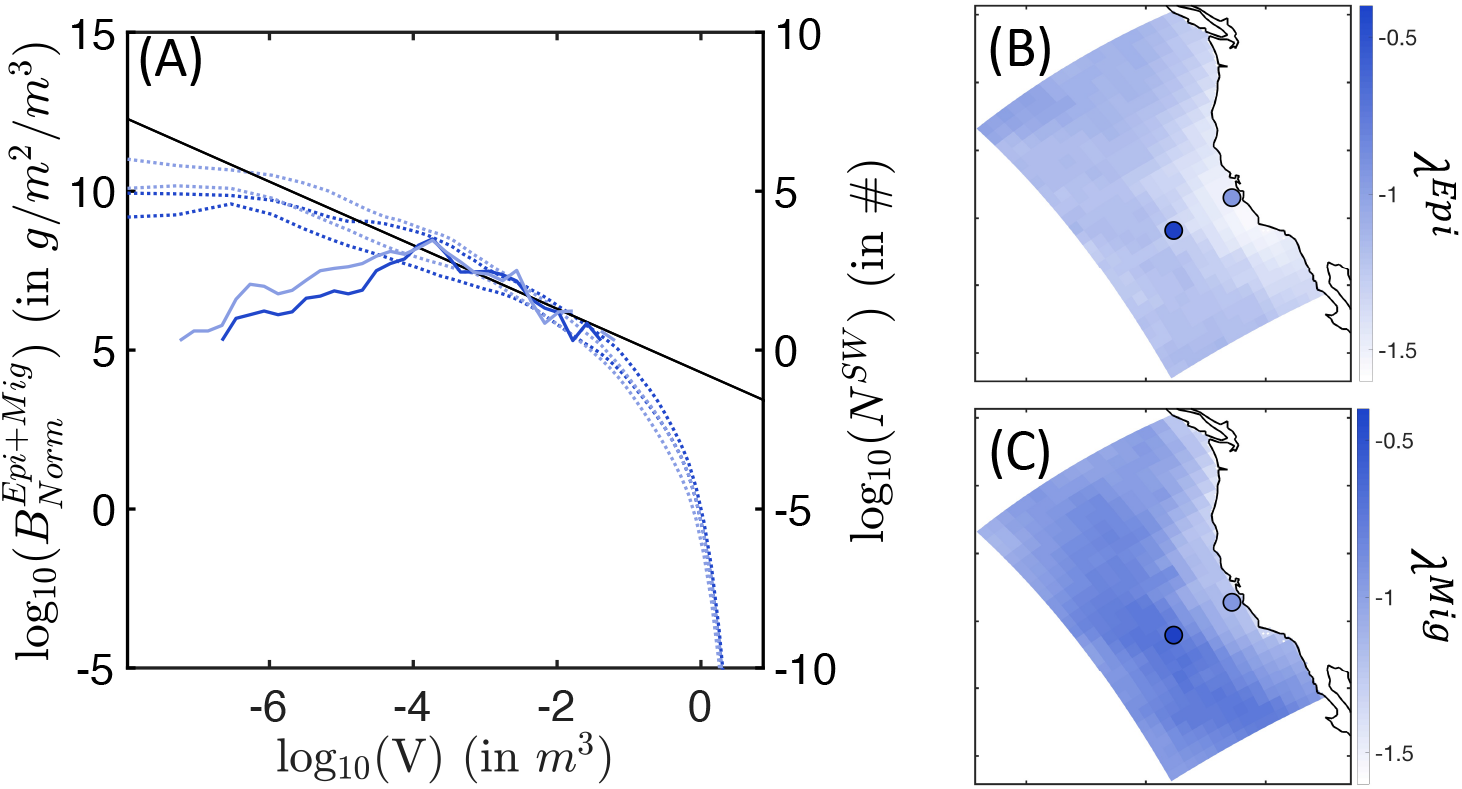
Biomass size spectra. (A) Envelop for the simulated normalised biomass sizespectra (dotted line) and observed biomass size-spectra (solid line), in a coastal (light blue) and an offshore (dark blue) cell. (B) Slopes λ^*Epi*^ of the local average size-spectra for the epipelagic community. (C) Slopes λ^*Mig*^ of the local average size-spectra for the migratory community. In panel (A), the black line shows the expected theoretical slope of λ = −1. In panels (B-C), the light/dark blue dots indicate the cells where coastal/offshore biomass spectra panel (A) come from.

For the epipelagic community, *λ^Epi^* is steeper close to the shore, and shallower offshore (see Fig. 8B). As the slope becomes less steep, the biomass of small fish decreases compared to the biomass of large individuals (see simulated spectra Fig. 8A). This relative decrease of small fish biomass in coastal relative to offshore waters agrees with observations, where fewer small individuals are found in surface trawls moving from the coast to the open ocean (solid lines in Fig. 8A). Note that observed size spectra break for sizes smaller than *V* = 10^−4^ *m*^3^. In this range, a systematic decrease in biomass abundance can be in part attributed to a decreasing selectivity of sampling gear to small individual sizes.

For the migratory community, simulated spectral slopes show a similar trend (see Fig. 8C), with generally shallower values. Thus, larger predators are relatively more abundant among migratory fish, as supported by the larger sizes reached by this community (see Fig.7).

### 3.3. Cross-shore biomass distributions

#### 3.3.1. A cross-shore size succession

Biomass size-spectra indicate a cross-shore variation of the relative abundance of larger and smaller fish in the California Current (Fig. 8). Figures 9A,B highlights this variation for small, medium, and large individuals along an average cross-shore section. For the epipelagic community, only small (0.03 < *L* < 0.3m) and medium-size (0.3 < *L* < 0.9m) fish survive in the domain (see species size distribution in Fig. 7A). Most small fish are found near the coast (Fig. 9A, 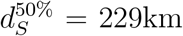), while medium-size fish are spread farther offshore 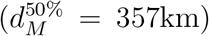. For the migratory community, in addition to small- and medium-size individuals, large predators (0.9 < *L* < 2m) survive. A similar pattern of increasingly spread-out biomass from small to large fish is also observed for this community (Fig. 9B, 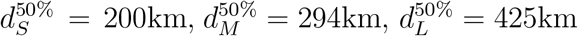).

**Figure 9:**
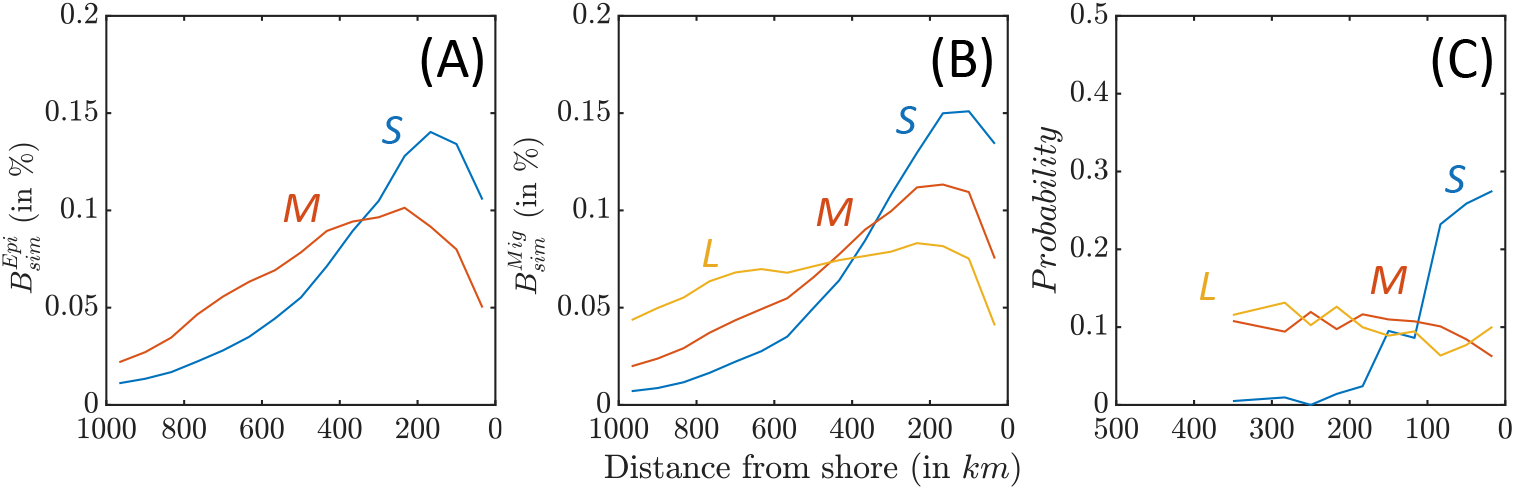
Biomass density along a cross-shore section, for small (blue), medium (red) and large size (yellow) fish. (A) Relative distribution of simulated epipelagic biomass distribution 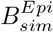. (B) Relative distribution of simulated migratory biomass distribution 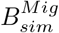. (C) Observed relative distribution from OBIS species occurrence data.

This cross-shore spread is comparable with the observed species distribution from the OBIS database (Fig. 9C, when all sample are considered, see Supplementary Information S8 for additional details). While small species (0.04 < *Lm* < 0.4m) cluster nearshore, medium (0.4 < *Lm* < 0.9m) and large (0.9*m* < *Lm*) species are more homogeneously distributed. Simulations and observations cannot be directly compared, because the OBIS database reports presence but not abundance. However, these patterns suggests higher occurrence of small fish in coastal water, while large fish tend to occupy the entire ecosystem.

Sensitivity tests suggest that the biomass spread for distinct size groups is mostly sensitive to the swimming parameter (*ADV*), and the predatorprey selectivity parameters (*k*_1,2_) (Fig. 6). For *ADV*, an increased swimming capability leads to offshore spread of migratory species, and vice versa. For *k*_1,2_, smaller predator-prey size ratios lead to a relative increase of biomass near the coast.

#### 3.3.2. Drivers of cross-shore succession

The best APECOSM-CC simulation is tuned to reproduce observed biomass densities. These densities are controlled by three main processes: (1) net or surplus production, modulated by environmental conditions; (2) current-driven redistribution from source to sink regions; (3) active swimming towards favorable conditions, i.e., from sink to source regions.

Figure 10A-C shows the difference between production and respiration (Δ_*META*_) for the epipelagic and migratory communities. Positive surplus production is observed in productive nearshore waters for smaller individuals in both communities (primary production shown in Supplementary Information S13). Moving offshore, surplus production rapidly decreases from positive to negative values, becoming a sink of biomass when mortality exceeds new production (Figs. 10A,B). Medium- and large-size species feeding on small fish tend to accumulate new biomass further offshore. This crossshore gradient in production is slightly stronger for the epipelagic community. The enhanced production nearshore and dissipation offshore are consistent throughout the CC region, except along the southern coast where production nearly equals mortality, leading to limited surplus production (Fig. 10C).

**Figure 10:**
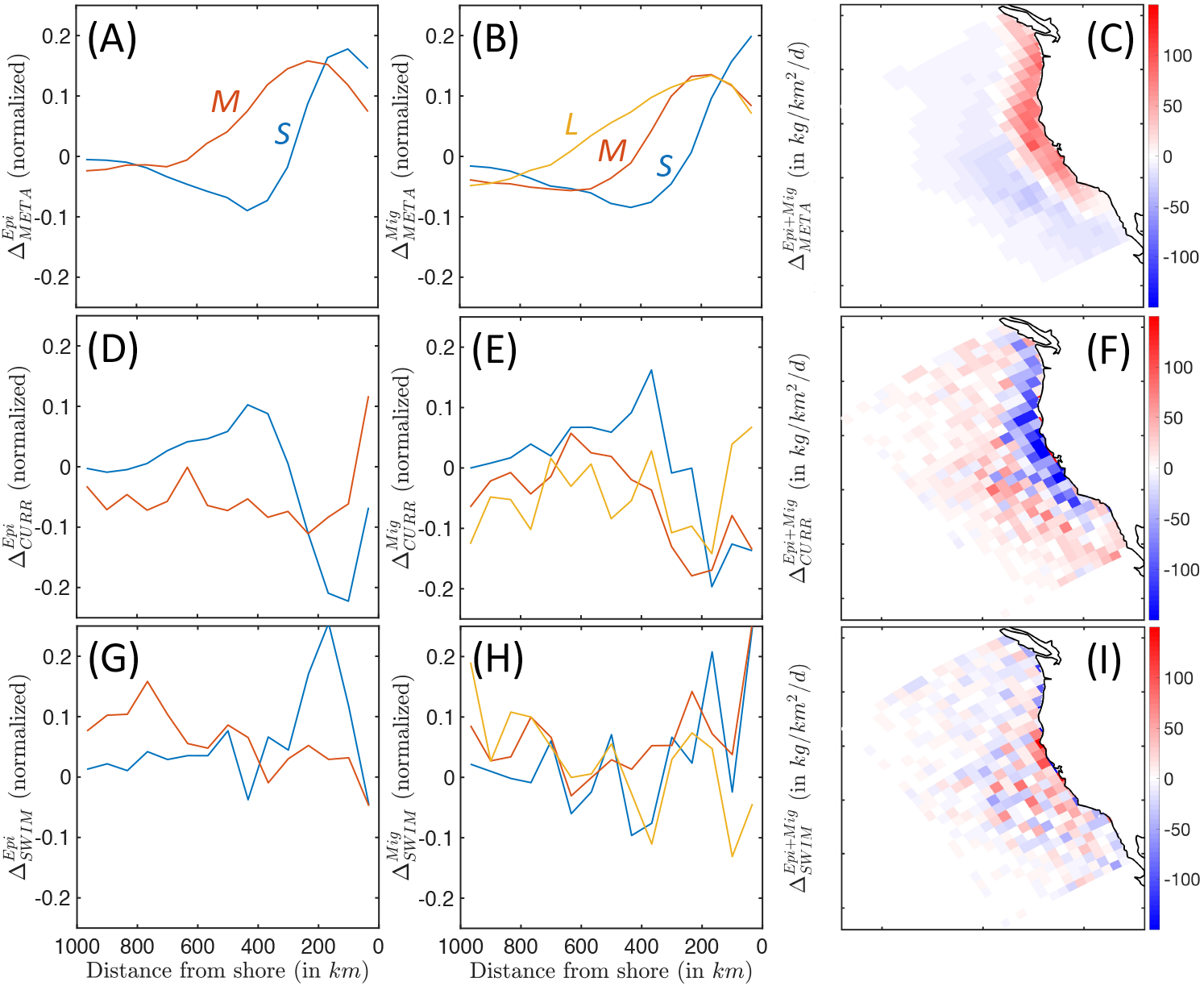
Biomass accumulation and dissipation along a cross-shore section, for small (blue), medium (red) and large (yellow) fish, reflecting food-web dynamics and spatial transport processes. (a-c) Surplus production for pelagic and migratory communities Δ_*META*_. (d-f) Accumulation and dissipation due to passive advection by currents for pelagic and migratory communities Δ_*CURR*_. (g-i) Accumulation and dissipation due to active swimming for pelagic and migratory communities Δ_*SWIM*_. The maps show averages binned in Δ*x* = 96km grid cells to smooth spatial variability caused by mesoscale eddies.

Currents act to redistribute biomass by advecting newly-produced biomass away from the coast, and accumulating it in a band between 300 and 800km, as revealed by Δ_*CURR*_ (see Figs. 10D-F). Most of this redistribution affects smaller individuals with limited swimming abilities, in both epipelagic and mesopelagic communities. As individuals grow larger, this transport is increasingly independent of the cross-shore distribution, and increasingly variable, suggesting an increasing interaction with mesoscale eddies, rather than continuous advection by the mean current (Figs. 10D,E). Since the bulk of biomass is found in smaller individuals, surplus biomass generated along the coast is on average advected offshore where it is mostly dissipated (see Figs. 10C,F).

In the model, swimming tends to counteract the effect of currents. Small fish in both communities swim on average towards the productive upwelling region, driving biomass accumulation along a band within 400km of the coast (see Δ_*SWIM*_, Figs. 10G,H). Swimming maintains the biomass of medium-sized migratory organisms along a similar coastal band (red line Fig. 10H), while larger migratory fish (yellow line Fig. 10H) and medium-sized epipelagic fish (red line Fig. 10G) tend to swim off-shore. To some extent, active movement against the mean current compensates the background transport offshore (compare the amplitude of passive advection Δ_*CURR*_ and active swimming Δ_*SWIM*_, Figs. 10F,I). However, active swimming also interacts with highly variable oceanic currents, such as eddies, fronts and filaments, diverting biomass from the coast to local features (as suggested by the increased patchiness in Fig. 10I).

In summary, while most new production is coastal, as individuals grow larger their main food source shifts offshore because of the transport by the mean surface current. Moreover, larger and larger individuals are increasingly attracted by local circulation features that divert them away from the most productive coastal regions. We argue that both effects contribute to explaining the cross-shore biomass distribution discussed in Section 3.3.1.

## 4. Discussion

### 4.1. Simulated biomass distribution

Our simulations reproduce observed relationships between fish biomass, temperature and primary productivity, with higher biomass in colder, high latitude and upwelling waters, and lower biomass in offshore oligotrophic waters. These gradient are best reproduced for the migratory community (Fig. 5A). Surface biomass densities are more homogeneously distributed over the domain, although with variations consistent with observations (Fig. 5B). However, the model underestimates the very large biomass densities in the most coastal eco-regions by up to a factor 10. This may be partly caused by the relatively low resolution of the model, which at 16 km poorly resolves the productive coastal band.

Observations of surface fish biomass density are also substantially more variable than model simulations. Surface-water trawl surveys are designed to monitor surface forage fish stocks (Zwolinski et al. (2012)), and therefore target biomass aggregations. These aggregations, which are shaped by fronts and other mesoscale features (Woodson and Litvin (2015), McGillicuddy Jr (2016)), are not fully resolved at the model resolution. Thus, the model should be considered a representation of more average conditions. Mid-water trawls sample the more homogeneous deep-scattering layer, and the reduced spatial variability of migrating mesopelagic fish is better captured by the model.

The biomass density for migrating mesopelagic fish is also underestimated by the model compared to the mid-water trawls, but matches other large-scale estimates for the region (e.g., 3.6 *gm*^−2^ in Lam and Pauly (2005)). Mesopelagic fish observations are associated with uncertainties, including trawl net avoidance and limited selectivity (Kaartvedt et al. (2012)). Some of these uncertainties are accounted for in the mid-water trawls data that we use in this study (Davison et al. (2013)). While additional fine-tuning of the model could help resolving this mismatch (see Fig. 6), we kept the optimal parameters for the final simulation to avoid generating other mismatches.

While the model underestimate mid-water trawl biomass, the ratio of pelagic to migratory biomass 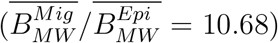 agrees with reconstructions for the California Current (Koslow and Davison (2016)), and other global estimates (Irigoien et al. (2014)). Furthermore, the model epipelagic biomass roughly matches observations, if we acknowledge the higher variability and patchiness of surface-water trawl data.

### 4.2. Parameter selection

Although mechanistic models often rely on fewer undetermined parameters than empirical models, some of these parameters remain poorly constrained. We conducted a sensitivity analysis to evaluate the importance of 11 of these parameters, and their effect on fish biomass distribution.

For predator-prey interactions, the optimal value for *C_FONC_* (see Table 1) indicates a specific volume clearance rate of 30.34 10^6^ per day (see Supplementary Information S2). We were not able to find measures for adult fish to confirm this number, but it is within the range measured from fish larvae and other lower trophic level organisms (Kiørboe (2011)). The parameter *CRISTAL_CRIT_* controls the intensity of feeding interactions, but is harder to compare to the literature. Tuning of this parameter mostly affects the relative biomass of epipelagic and migratory communities (see Fig. 6).

Parameters that determine the shape of the prey size selectivity (*k*_1,2_) influence the predator-prey mass ratio *PPMR* and the ability of predators to feed on a wider range of prey sizes (*σ*). Here, the best values for *k*_1,2_ (Table 1), correspond to a *PPMR* = 4861, and a selectivity width of approximately one order of magnitude (*σ* = 1.13, see also Supplementary Information S4). The estimated *PPMR* and the variability of the selectivity are well within the range of expected values (Barnes et al. (2010), Reum et al. (2019)). Consistently with theories that suggest that *PPMR* and *σ* have a strong effect on trophic efficiency (Brown et al. (2004), Eddy et al. (2020)), these parameters most affect the model solution, with longer food-webs (small *PPMR*) leading to more biomass accumulation (see Fig. 6). Moreover, both parameters influence the maximum sizes reached by communities (refer to the impact of *k*_1,2_ on 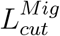 in Fig. 6).

The external mortality influences the total biomass accumulation, and the relative biomass of different size groups (Fig. 6). In particular, external mortality triggers a trophic cascade: as mortality increases, the biomass of larger individuals drops, favoring accumulation of small individuals. The optimum mortality value (see Table 1) represents a daily mortality at any size class *L* of 0.0078 *d*^−1^ when at maximum schooling density. The biomass in size classes with high densities will be lost with a time scale of approximately 4 months, while more diffuse size classes will decrease more slowly. Note that this mortality term includes any density-dependent extrinsic mortality. Thus, removal by fishing, which is not explicitly included in our simulations, is partly resolved by this term.

Coupling of APECOSM-CC to ROMS-BEC appears to be weakly sensitive to the size range of diatoms, zooplankton, and particulate organic carbon. Instead, the spatial distribution of these lower trophic levels is more important, because it determines gradients in suitable habitats. For example, the cross-shore biomass distribution for small fish (*B_S_* in Figs. 9A,B) closely matches the distribution of their food (Supplementary Information S13). Compared to observations from the MAREDAT database (Moriarty and O’Brien (2013)), ROMS-BEC simulations underestimate zooplankton biomass nearshore, and overestimate it offshore (see Kessouri et al. (2020) and Supplementary Information S13). We suggest that this bias in lower trophic levels is partly responsible for an excessive spread of fish biomass offshore in our simulations.

### 4.3. Emergent features

The model is able to capture a series of emergent properties of the observed biomass distribution in the California Current.

First, simulations show higher biomass of small migratory mesopelagic fish (see Fig. 4A), consistent with observations (Irigoien et al. (2014), Koslow and Davison (2016), Davison et al. (2013)). Since in the model surface epipelagic and deep-diving migratory communities share the same parameterizations, this property emerges from differences in their habitat. We suggest that differences in temperature, and the resulting metabolic rates, are behind these patterns. Surface epipelagic fish experience comparatively warmer conditions throughout the day, while migratory organisms experience colder waters during the day when they reside in the mesopelagic zone. Reduced respiration rates in cold waters would favor biomass accumulation of migratory mesopelagic fish. Note that the resident mesopelagic community that does not migrate is not modeled here, unlike in other applications of APECOSM (Maury (2010)). When included in the simulations, we are unable to model the three coexisting communities. Inclusion of this community will be explored in future studies.

Second, epipelagic fish tend to be smaller than migratory fish (see Fig. 7). While observed and simulated representative sizes are not perfectly comparable, the model reproduces a clear size difference between surface and migratory communities that agrees with observations. We suggest that the higher migratory biomass and lower metabolism in deep waters allow the development of more trophic levels in the migrating community, ultimately supporting larger species and individuals (Guiet et al. (2016)). Note that the maximum size reached by migratory predators 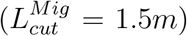 is below the maximum size allowed by the model (2m), while observations show the occurrence of larger species 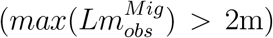. Most large deep-diving predators develop thermoregulation apparatuses that increase feeding efficiency and assimilation in deep waters. These adaptations may contribute to the increased asymptotic size of migratory species, but are absent in the model. Fine-tuning or new parameterizations could help reduce this bias (see Fig. 6).

Third, an ubiquitous feature of marine ecosystems is the regularity of the biomass density distribution, which follows a power-law as a function of individuals size, at least within selected size ranges (Brown et al. (2004), Hatton et al. (2021)). For the biomass density as a function of the individual structural volume (*V* = (*δL*)^3^), the slope of this power-law is empirically and theoretically shown to be close to λ ≈ – 1 (Brown et al. (2004), Rossberg (2012), Sprules and Barth (2015)). APECOSM-CC reproduces this emergent slope, with 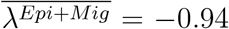, and regional variations and inter-community differences (see Fig. 8). Simulated variations around this mean value are large, since slopes as steep as −1.57 for the epipelagic community in the core of the upwelling, and as shallow as −0.71 offshore for the migratory community are simulated. The model suggests comparatively more small individuals nearshore, and fewer offshore. We argue that this shift is due to biomass redistribution between source and sink regions. Note that fishing has been shown to impact the slope of fish size spectra (Shin and Cury (2004)), an effect not explicitly included in our simulations.

### 4.4. Cross-shore biomass spread

The model reveals an increase in the cross-shore biomass spread for individuals of increasing size (see Figs. 9A,B). The pattern is comparable to observations from the OBIS database (Figs. 9C), showing that smaller species are sampled more frequently along the coast, while larger species are sampled more uniformly across the region. Importantly, this cross-shore pattern extends to higher trophic levels a succession already documented for phytoplankton and zooplankton (Messié and Chavez (2017), Keister et al. (2009)).

This cross-shore distribution emerges in the model from surplus biomass production for small organisms near the coast, where lower trophic levels are more abundant. This excess production, likely trapped in the surface Ekman layer and mesoscale features, is then advected offshore, as observed for zoo-plankton in the California Current (Keister et al. (2009)). Small fish biomass accumulates while being transported offshore, in turn feeding increasingly large predators. Unlike zooplankton, fish are able to swim towards more favorable conditions. Therefore we would expect all organisms to remain in the core of the productive upwelling region. However, (1) only individuals longer than approximately 0.13m can swim against the average surface currents, while the rest is slowly advected offshore; (2) intense mesoscale currents and eddies lead to patchy biomass distributions, with organisms clustering around mesoscale eddies seeded with coastal biomass. Increasingly large predators are attracted by these high-biomass “oases”, limiting their ability to constantly swim towards the coast (Figs. 10G,H). Increased biomass aggregations along eddies and fronts are further expected to affect the productivity of higher trophic levels (Woodson and Litvin (2015)).

These cross-shore patterns depend on parameters that control predatorprey interactions. Shorter food-webs (i.e., larger PPMR) lead to enhanced coupling with lower trophic levels, and a biomass distribution that increases more rapidly towards the coast (see 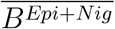 and *d*_50%_ for various values of *k*_1,2_, Fig. 6). We suggest that with a larger *PPMR*, the distribution of primary consumers matches more closely the higher primary production near the coast, so that secondary consumers will also cluster onshore following their prey. Note that in our simulations the biomass of smaller mid-trophic levels is distributed more broadly offshore than suggested by the OBIS database. As shown by Figure 9, small organisms dominate within 400km from the coast (blue lines in panels A and B) while in observation they dominate within 100km from the coast (blue line in panel C). This bias can be partly attributed to biases in lower trophic level biomass in ROMS-BEC (see comparison with MAREDAT data, Moriarty and O’Brien (2013), in Supplementary Information S13).

## 5 Concluding remarks

In this paper, to test the influence of movement on the structure of fish communities, we implement APECOSM-CC, a mechanistic model of the fish biomass flow in the California Current. We also process a diverse set of observations from this densely sampled ecosystem, to describe the main characteristics of the epipelagic and migratory fish biomass distribution, and provide a series of observational constrains to optimize uncertain parameters and evaluate the model. The model is calibrated to reproduce spatial patterns and magnitude of bulk fish biomass for epipelagic and mesopelagic migratory communities, but it also reproduces a series of emergent properties suggested by observations. These include: (1) larger and more abundant migrating predators relative to epipelagic fish; and (2) a cross-shore succession in the size of fish, with smaller individuals clustered along the coast, and larger individuals increasingly spread offshore.

Our study is the first to couple a mechanistic, size-structured food web model to a dynamically evolving representation of ocean currents that resolves mesoscale eddies. In the very dynamical California Current it indicates three essential effects of transport and movement: (1) the importance of the surface mean current in advecting small individuals offshore, setting the cross-shore size structure of fish communities; (2) the role of active swimming in counterbalancing this transport; and (3) the importance of eddies in limiting the effectiveness of swimming and contributing to a more homogeneous distribution of predators across the domain.

Because of its mechanistic formulation and realistic forcing, APECOSM-CC represents an ideal tool to continue investigating fundamental ecological processes that regulate marine fish communities in a dynamical environment, their interactions with fine-scale ocean currents, and the effects of environmental variability, from natural fluctuations, to climate change, fishing, and other human activities.

## 6. Acknowledgment

This research was supported by California Ocean Protection Council grant C0100400. Computational resources were provided by the Extreme Science and Engineering Discovery Environment (XSEDE) through allocation TG-OCE170017, and by the super-computer Hoffman2 at the University of California Los Angeles, at the Institute for Digital Research and Education (IDRE, UCLA). Daniele Bianchi acknowledges additional funding from the Alfred P. Sloan Foundation.

