## Supplementary Materials for "Movement shapes the structure of fish communities along a cross-shore section in the California Current"

### S1: Parameters of APECOSM

There are 48 parameters and constants in APECOSM, for the food-web parameters and the spatial dynamics (Tables 1 and 2). Many of these parameters are well constrained by the literature. The parameters of the DEB theory are based on a meta-analysis of observations of fish life-histories (Kooijman and Lika (2014)) as well as fine tuning to account for realistic growth (Guiet et al. (2016)). The sensitivity to temperature and other environmental variables, and the implementation of movement, have been previously described (Maury (2010), Guiet et al. (2016)).

Only 7 parameters describing predator-prey interactions, and 6 parameters controlling coupling with ROM-BEC remain to be specified. Among these, the size dependence of half saturation constant ($C_{FONC\ W\ DEP}$) is prescribed as 1/3, based on mechanistic assumptions for fish feeding. The schooling parameter which controls the strength of the density-dependence of schooling ($CRISTAL_{SLOPE}$) is set to 2. We estimate the remaining 11 parameters using an ensemble of simulations.

Table 1: Food-web parameters in APECOSM-CC. For undetermined parameters (bold fonts) the values provided correspond to the best ensemble selected.

| Parameter | Name | Unit | Value | |
|---|---|---|---|---|
| $[L_{min},L_{max}]$ | Minimum/Maximum size of higher trophic levels | $m$ | $[10^{-3}$,2] | |
| $[Lm_{min},Lm_{max}]$ | Minimum/Maximum species size | $m$ | $[10^{-3}$,2] | – |
| $n_{bins}$ | Number of size bins | – | 50 | |
| $n_{spec}$ | Number of species | – | 6 | |
| $a_{pAm}$ | Assimilation rate coefficient | $J/m^3/d$ | $31.25\ 10^6$ | |
| $a_{Em}$ | Reserve density coefficient | $J/m^4$ | $312.5\ 10^8$ | |
| $p_M$ | Maintenance rate | $J/m^3/d$ | $25\ 10^6$ | |
| $E_g$ | Volume specific cost of growth | $J/m^3$ | $5691\ 10^6$ | |
| $K$ | Energy fraction allocated to growth and maintenance | – | 0.8 | |
| $K_r$ | Energy fraction of gonads turned into eggs | – | 0.95 | |
| $e_m$ | Assimilation efficiency | – | 0.8 | |
| $h_a$ | Aging acceleration | $d^{-2}$ | $10^{-6}$ | |
| $\Psi$ | Energy content of biomass | $J/g$ | 4552 | |
| $d$ | Density of biomass | $g/m^3$ | $10^6$ | |
| $F_{egg}$ | Fraction of spawned eggs not fertilized | – | 0.2 | |
| $\varphi$ | Sex ratio | – | 0.5 | |
| $\alpha_p$ | Puberty structural volume coefficient | – | 0.125 | |
| $\delta$ | Structural volume/length factor | – | 0.2275 | |
| $T_A$ | Mean Arrhenius temperature | $^oK$ | 8000 | |
| $T_{ref}$ | Reference temperature | $^oK$ | 293.15 | |
| $Q$ | Scaling exponent of the attack rate | – | 1. | |
| $P$ | Scaling exponent of the handling time | – | 1./3. | |
| $\mathbf{C_{FONC}}$ | Half saturation constant | $J/m^3$ | **1.03** | |
| $C_{FONC\ W\ DEP}$ | Weight dependence of half saturation constant | 1/3 | – | |
| $\mathbf{CRISTAL_{CRIT}}$ | Schooling intercept | $(J/m^3)$ | **133** | |
| $CRISTAL_{SLOPE}$ | Schooling transition slope | – | 2 | |
| $\mathbf{k_1}$ | Factor 1 for predator/prey selectivity | – | **1.16** | |
| $\mathbf{k_2}$ | Factor 2 for predator/prey selectivity | – | **2.05** | |
| $\mathbf{M}$ | External mortality rate | $d^{-1}$ | **0.0078** | |
| $\mathbf{L_{min}^{Diat}}$ | Minimum size of diatoms | $m$ | $\mathbf{5.2\ 10^{-6}}$ | |
| $\mathbf{L_{max}^{Diat}}$ | Maximum size of diatoms | $m$ | $\mathbf{1.4\ 10^{-4}}$ | |
| $\mathbf{L_{min}^{Zoo}}$ | Minimum size of meso-zooplankton | $m$ | $\mathbf{1.6\ 10^{-4}}$ | |
| $\mathbf{L_{max}^{Zoo}}$ | Maximum size of meso-zooplankton | $m$ | $\mathbf{1.1\ 10^{-2}}$ | |
| $\mathbf{L_{min}^{POC}}$ | Minimum size of POC | $m$ | $\mathbf{1.7\ 10^{-4}}$ | |
| $\mathbf{L_{max}^{POC}}$ | Maximum size of POC | $m$ | $\mathbf{1.4\ 10^{-2}}$ | |

Table 2: Dynamic parameters in APECOSM-CC.

| Parameter | Name | Unit | Value |
|---|---|---|---|
| $\delta t$ | Advection/diffusion time step | $day$ | 0.139 |
| $DIFF_{PHYS}$ | Diffusivity of an inert particle | $m^2/day$ | $17.\ 10^6$ |
| $DIFF$ | Diffusivity of an organism of length $L_{max}$ | $m^2/day$ | $100.\ 10^6$ |
| $ADV$ | Advection of swimming individual for 1m organisms | $m/day$ | $45.\ 10^3$ |
| $K_{ADV}$ | Half saturation constant of advection | $m^{-1}$ | $2.5\ 10^{-7}$ |
| $DzPHY$ | Vertical diffusivity coefficient | $m^2/day$ | 0.35 |
| $DIFFz$ | Vertical diffusion for 1m organism (per community) | $m^2/day$ | $[52.\ 10^3, 45.\ 10^3]$ |
| $ADVz$ | Vertical advection for 1m organism (per community) | $m/day$ | $[45.\ 10^3, 45.\ 10^3]$ |
| $\sigma_{TCOR}$ | Standard deviation of preferred temperature (per community) | $^oK$ | $[0.1, 0.]$ |
| $O_2^{Lim}$ | Threshold of $O_2$ (per community) | $\mu mol/L$ | $[10^{-4}, 0]$ |
| $O_2^{Resp}$ | Flatness of sigmoid response of O2 (per community) | $\mu mol/L$ | $[10^5, 0, 0]$ |
| $OPT_{LIGHT}$ | Optimal light (per community) | $W/m^2$ | $[10^2, 10^{-3}]$ |
| $\sigma_{LIGHT}$ | Standard error of light (per community) | $W/m^2$ | $[1.7\ 10^2, 8.\ 10^{-3}]$ |

### S2: Prey encounter rate and $C_{FONC}$

Following the description of APECOSM in Maury and Poggiale (2013), a functional responses $f$ describes the intake rate of a consumer as a function of prey density $p$ (in $J/m^3$ in APECOSM). The model adopts a type-II functional response that describes this intake as a function of handling time $h$ (in $d/J$), the time a predator needs to ingest a prey, and encounter/attack rate $e$, the rate of prey captured per time unit (in $m^3/d$):

$$f = \frac{p}{\frac{1}{he} + p} \tag{1}$$

In APECOSM, the encounter/attack rate is proportional to the length of fish ($L = V^{1/3}/\delta$) and the surface of the capturing apparatus ($V^{2/3}$), such that $e = K_1 V^{1/3} V^{2/3} = K_1\ V$ ($K_1$ the volume specific clearance rate in $d^{-1}$). The handling time is inversely proportional to the assimilation rate, such that $h = 1/a_{pAm}\ V_m^{-1/3} V^{-2/3}$ (in $d/J$):

$$f = \frac{p}{\frac{1}{he} + p} = \frac{p}{p + (a_{pAm}\ V_m^{1/3} V^{2/3})/(K_1 V)} = \frac{p}{p + C_{FONC} V_m^{1/3} / V^{C_{FONC\_W\_DEP}}} \tag{2}$$

where $C\_FONC = a_{pAm}/K_1$ and $C\_FONC\_W\_DEP = 1/3$.

Based on estimates of the clearance rate (Hartvig et al. (2011), Kiørboe (2011)), we select the undetermined parameter $K_1 \in [10^6, 5.\ 10^8]$. The corresponding half saturation constant is $C\_FONC = a_{pAm}/K_1 \in [0.0625, 31.25]$.

### S3: Schooling and $\mathbf{CRISTAL_{CRIT}}$

In APECOSM, the prey density $\xi_V$ (in $J/m^3/m^3$) at structural volume $V$ (in $m^3$) available to predators depends on a schooling probability $ps$ which is a function of this same prey density $\xi_V$:

$$ps_{V,\xi_V} = \frac{(V\xi_V)^{CRISTAL_{SLOPE}}}{(V\xi_V)^{CRISTAL_{SLOPE}} + (CRISTAL_{CRIT})^{CRISTAL_{SLOPE}}} \quad (3)$$

where schooling transition slope $CRISTAL_{SLOPE}$ and schooling intercept $CRISTAL_{CRIT}$ are parameters that set the shape of the distribution for $ps$. We select $CRISTAL_{SLOPE} = 2$ and estimate $CRISTAL_{CRIT}$ such that at global maximum and minimum biomass at low trophic levels $B^{LTL}$ ($B^{Diat}$, $B^{Zoo}$ and $B^{POC}$), schooling $ps^{LTL}$ is respectively above 0.25 and below 0.75, meaning at least 25% and up to 75% of low trophic level biomass is accessible.

To link biomass at low trophic level provided in the forcing, $B^{LTL}$, with low trophic level prey density $\xi_V^{LTL}$, we assume the biomass density as a function of structural volume a power law of slope $-1$, $\xi_V^{LTL} = \xi_0 V^{-1}$ (see S6), such that:

$$B^{LTL} = \int_{V_{min}^{LTL}}^{V_{max}^{LTL}} \xi_0 V^{-1} dV \quad (4)$$

Equation 4 is solved to estimate $\xi_0$ such that $\xi_V^{LTL}$ writes:

$$\xi_V^{LTL} = \frac{B^{LTL}}{ln(V_{max}^{LTL}/V_{min}^{LTL})} V^{-1} = \frac{B^{LTL}}{3\ ln(L_{max}^{LTL}/L_{min}^{LTL})} V^{-1} \quad (5)$$

assuming $V = (\delta L)^3$. Then, the schooling probability at low trophic level (eq. 3) is:

$$ps^{LTL} = \frac{1}{1 + (3\ ln(L_{max}^{LTL}/L_{min}^{LTL})\ CRISTAL_{CRIT}/B^{LTL})^{CRISTAL_{SLOPE}}} \quad (6)$$

The size range of low trophic level communities $[L_{min}^{LTL}, L_{max}^{LTL}]$ is set to vary (see S6). Once sets of size ranges are chosen, we solve equation 6 for $ps^{LTL} = 0.25$ and $max(B^{LTL}) = 39400,\ 3645,\ 8590\ J/m^3$ for respectively diatoms, zooplankton and particulate organic carbon, for $ps^{LTL} = 0.75$ and $min(B^{LTL}) = max(B^{LTL})/1000$.

With this approach the $CRISTAL_{CRIT}$ varies on the range $[0.15, 9880]$ $J/m^3$, which corresponds to $[0.25, 3.8\ 10^{-6}]$ $g/m^3$.

### S4: Prey selectivity and $(\mathbf{k_1, k_2})$

Following Maury and Poggiale (2013), the selectivity $s_{u,v}$ of a predator of mass $u$ on a prey of mass $v$ is determined by a combination of two sigmoid functions (see Fig. 1A):

$$s_{u,v} = \frac{1}{1 + e^{(\alpha_1(\rho_1 - (u/v)^{1/3}))}} \left(1 - \frac{1}{1 + e^{(\alpha_2(\rho_2 - (u/v)^{1/3}))}}\right) \tag{7}$$

for $\alpha_{1,2}$ controlling the slope and $(\rho_{1,2} - (u/v)^{1/3})$ controlling the position of the inflection point for each sigmoid function. In order to test the influence of this selectivity on the biomass flow in the food web, we allow various shapes multiplying $(u/v)^{1/3}$ by $k_1$ and $\alpha_{1,2}$ by $k_2$. Multiplying by $k_1$ leads to a shift of the selectivity function, i.e. and increase or decrease of the ratio between the mass of preys and predators. Multiplying by $k_2$ influences slopes leading to a flattening or narrowing of the selectivity function (see Fig. 1A).

$$s_{u,v} = \frac{1}{1 + e^{(k_2 \ \alpha_1(\rho_1 - k_1 \ (u/v)^{1/3}))}} \left(1 - \frac{1}{1 + e^{(k_2 \ \alpha_2(\rho_2 - k_1 \ (u/v)^{1/3}))}}\right) \tag{8}$$

The multiplication of the selectivity function $s_{u,v}$ by an idealized prey biomass density distribution $\xi = \xi_0 V^{-1}$ allows the identification of a realised predator prey mass ratio $PPMR$ and realised selectivity width $\sigma$ (Fig. 1B). The predator prey mass ratio is estimated as the ratio $PPMR = u_0/v_{50}$ between a reference predator mass, $u_0 = 1.87 \ 10^{-4}\text{m}^3$ corresponding to a 0.25m long predator, and the volume $v_{50}$ at mid-point of the distribution $S_{u,v} = s_{u,v}\xi$. The selectivity width is taken as the ratio between the 10th and 90th percentiles on a logarithmic scale $\sigma = log_{10}(v_{90}/v_{10})$ (Fig. 1B).

With these definitions, varying $k_{1,2} \in [1, 3]$, the $PPMR$ values are between 245 and 19260 for a 0.25m long predator while the width $\sigma$ varies between 1 and 1.7 orders of magnitudes.

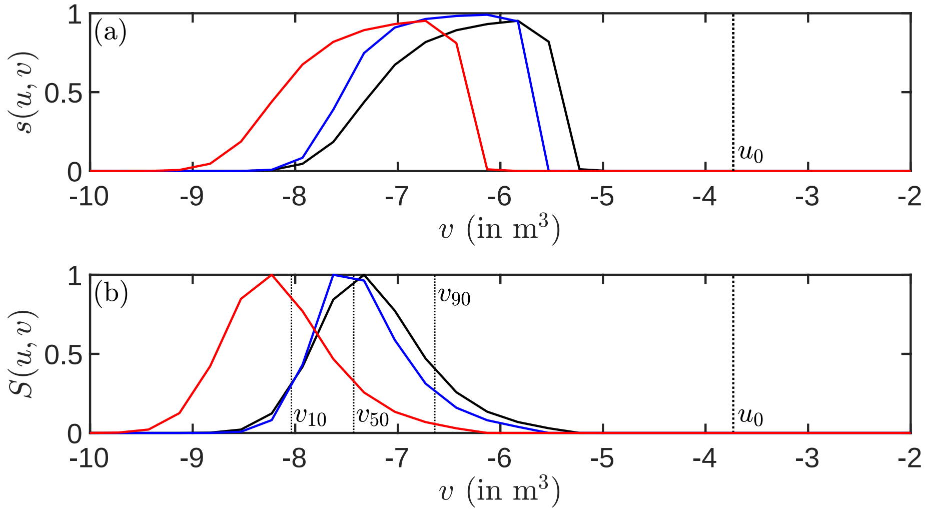


Figure 1: Prey selectivity function: (A) examples of selectivity distribution $s_{u,v}$ for a predator size $u = u_0$; (B) examples of realised selectivity $S_{u,v}$ for a predator size $u = u_0$. The black curves illustrate a reference distribution that is shifted when varying $k_1$ (red curves), that is narrowed when varying $k_2$ (blue curves).

### S5: Background mortality M

Multiple source of mortality are included in APECOSM, including predation mortality, ageing and starvation. In addition, we parameterize an external mortality $Z$ which depends on fish aggregation as described with schooling probability $ps_{V,\xi_V}$ (eq. 3):

$$Z_{V,\xi_V} = M\ ps_{V,\xi_V} \xi_V \tag{9}$$

for individuals of structural volume $V$ and a biomass density $\xi_V$, with $M$ (in $d^{-1}$). We estimate the parameter $M$ such that 100% of individuals in the low trophic level compartments die within a week to within 6 month due to disease, predation by unrepresented functional groups. This is equivalent to $M\ ps^{LTL} \in [0.0056, 0.14]\ d^{-1}$.

With these criteria, and for $CRISTAL_{CRIT} \in [0.15, 9880]\ J/m^3$, the mortality constant $M$ varies between $[0.0055, 196]$.

### S6: Low trophic level size ranges

Biomass at lower trophic levels $B^{LTL}$ ($B^{Diat}$, $B^{Zoo}$ and $B^{POC}$, in $J/m^3$) is a key forcing in APECOSM. Because of the size dependent predation, this biomass is distributed as a function of individuals/particles sizes $V$ assuming a power law of slope $-1$:

$$\xi_V^{LTL} = \xi_0 V^{-1} \tag{10}$$

where the intercept $\xi_0$ depends on the total biomass for the trophic level group $B^{LTL}$:

$$B^{LTL} = \int_{V_{min}^{LTL}}^{V_{max}^{LTL}} \xi_0 V^{-1} dV \tag{11}$$

that once solved to estimate $\xi_0$ gives the distribution

$$\xi_V^{LTL} = \frac{B^{LTL}}{ln(V_{max}^{LTL}/V_{min}^{LTL})} V^{-1} = \frac{B^{LTL}}{3\ ln(L_{max}^{LTL}/L_{min}^{LTL})} V^{-1} \tag{12}$$

for $[V_{min}^{LTL}, V_{max}^{LTL}]$, or $[L_{min}^{LTL}, L_{max}^{LTL}]$, common size ranges for each low trophic level group ($Diat$, $Zoo$ or $POC$).

The selection of the size ranges influences the intercept of the low trophic level biomass density spectrum, and ultimately the prey available to upper trophic level predators. We allow the upper/lower size for each low trophic level group to vary around reference values $Lref_{min}^{Diat} = 10^{-5}$, $Lref_{max}^{Diat} = 10^{-4}$, $Lref_{min}^{Zoo} = 2.\ 10^{-5}$, $Lref_{max}^{Zoo} = 2.\ 10^{-3}$, $Lref_{min}^{POC} = 10^{-4}$ and $Lref_{max}^{POC} = 5.\ 10^{-3}$ (in $m$).

### S7: Properties of SW trawls

We use the Surface-Water (SW) trawls to estimate the observed biomass density distribution of epipelagic species, $B_{obs}^{Epi}$. For comparison with the model, we consider samples taken at night, since most SW trawls are nocturnal, and sum the weight of contributing species before normalizing by the volume of water filtered by the trawl. SW trawls include multiple functional groups. Here we focus on fish species, especially pelagic species classified in Fishbase as belonging to any of the following categories, *pelagic*, *pelagic-oceanic*, *pelagic-neritic* or *bathypelagic*. Figure 2A shows the biomass density distribution $B_{obs}^{Pel}$ of all trawls where pelagic species have been sampled ($\overline{B_{obs}}^{Pel} = 0.94\ g/m^2$). Figure 2B-D breaks this distribution into the functional groups simulated with APECOSM: $B_{obs}^{Epi}$, for *pelagic*, *pelagic-oceanic* and *pelagic-neritic* Fishbase groups with a diving depth above 150m; $B_{obs}^{Meso+Mig}$ for *pelagic*, *pelagic-oceanic*, *pelagic-neritic* Fishbase groups with a diving depth below 150m, and *bathypelagic*; $B_{obs}^{Meso}$ for the *bathypelagic* Fishbase group. The distributions figures 2 range up to 6 orders of magnitude, we suggest because of the patchiness of the fish biomass distribution. In average, the epipeplagic biomass density $\overline{B_{obs}}^{Epi} = 1.1\ g/m^2$ is higher than the mesopelagic and migratory biomass density $\overline{B_{obs}}^{Meso+Mig} = 0.31\ g/m^2$, contradicting the expectation of about one order of magnitude more mesopelagic biomass than epipelagic biomass (Koslow and Davison (2016), Irigoien et al. (2014)). Although for these night trawls migratory mesopelagic fish come at the surface, the mesopelagic biomass density is a small proportion (see Figure 2D and $\overline{B_{obs}}^{Meso} = 1.8\ 10^{-2}\ g/m^2$). Sampling the upper 30m of the ocean may not reach the mesopelagic migrators. Therefore, we only compare SW trawls with simulated epipelagic biomass densities.

The identification of species contributing to each trawl as well as the weight sampled per species per trawl $w_s$ allows the estimation of the average species size per trawl $\overline{Lm}_{obs} = \sum Lm_s w_s / \sum w_s$ ($Lm_s$ the asymptotic length of a species as provided in Fishbase). Keeping the previously described criteria in order to identify sub-groups in trawls, we determine the asymptotic size distribution for each community. Figure 3A-C shows clear differences between communities as revealed by the median mean asymptotic lengths $\overline{Lm}_{obs}$, 0.4m for epipelagic species, 0.91m for diving epipelagic predators predators and 0.12m for mesopelagic migrators.

Finally, for a subset of the SW trawls, individuals are measured in addition of being weighted. We use these 22210 length measurements to test the expectation of an abundance distribution that follows a power-law (Andersen et al. (2015)). From 0.25 to 4m the biomass distribution matches expectations, with a slope $\lambda = -3.05$. Below 0.25m the abundance distribution diverges from the power-law distribution, we expect SW trawls to slightly under-sample mid-trophic level epipelagic biomass (Fig. 4).

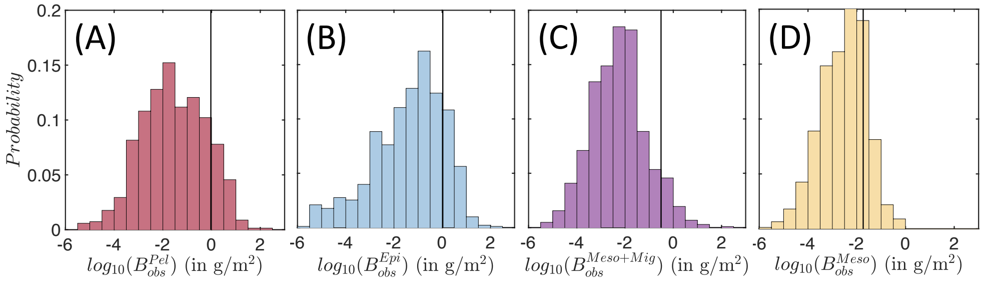


Figure 2: Biomass density distribution $B_{obs}$ in SW trawls: (A) all pelagic fish; (B) epipelagic fish; (C) mesopelagic and migratory pelagic fish; (D) mesopelagic fish. In each panel the black vertical line indicates the mean biomass density.

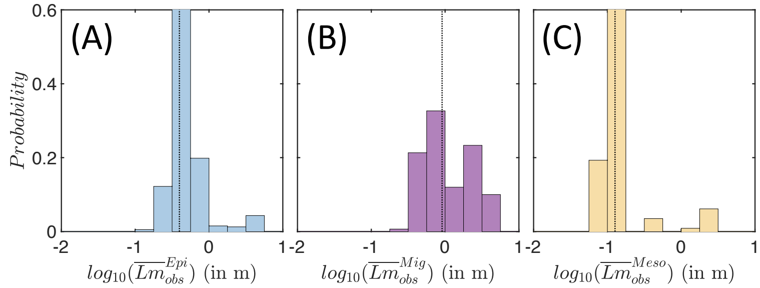


Figure 3: Mean species size distribution $\overline{Lm}_{obs}$ in SW trawls: (A) epipelagic fish; (B) migratory pelagic fish; (C) mesopelagic fish. In each panel the black doted vertical line indicates the median asymptotic length.

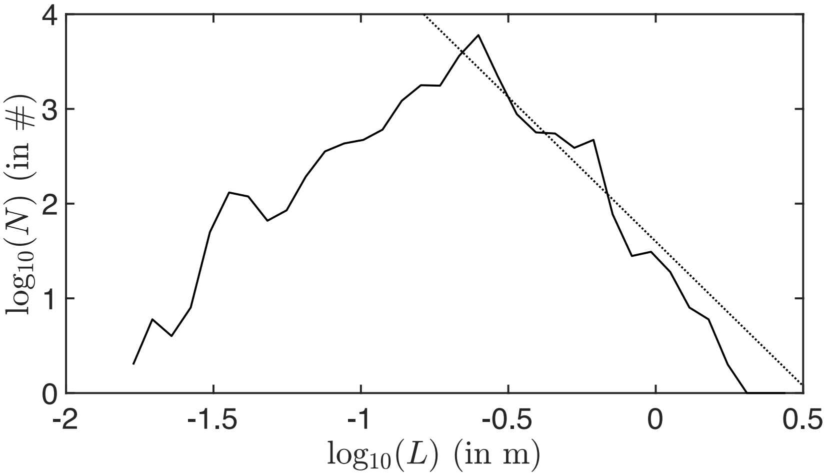


Figure 4: Abundance distribution as a function of measured individual's size. The dashed line illustrates a power law of slope $\lambda = -3.05$ that matches the abundance distribution of larger individuals.

### S8: Properties of OBIS data

OBIS data inform the spatio-temporal occurrence of species at distinct life stages, at distinct depths (OBIS). Inside our domain of the California Current, it provides observation for 1369 species. We match the list of species with Fishbase (Froese and Pauly (2016)) in order to separately select epipelagic or mesopelagic fish samples, i.e. respectively species labeled as *pelagic*, *pelagic-oceanic*, *pelagic-neritic* or *bathypelagic*. As in the previous section we also use depth ranges provided in Fishbase to differentiate surface epipelagic species from vertically migrating predators. This leads to $n_{obs}^{Epi} = 4088$ observation of surface epipelagic fish, $n_{obs}^{Mig} = 10279$ observation of vertically migrating epipelagic fish and $n_{obs}^{Meso} = 11638$ observation of mesopelagic fish. For each sample we determine the asymptotic length $Lm$ from the value prescribed in Fishbase (Froese and Pauly (2016)) for the corresponding species. Figure 5 merges the spatial occurrence of samples belonging to each of the aforementioned fish communities with the asymptotic lengths in order to reveal the cross-shore probability of occurrence per species of increasing size. To determine this probability, we first estimate the proportion $R^i$ of individuals from small ($0.04 < Lm < 0.4$m), medium ($0.4 < Lm < 0.9$m) and large ($0.9 < Lm < 2$m) species in regular distance bins $i$ ($dx_i = 33$km) from the nearest shore, $R^i_{S,M,L} = n^i_{S,M,L}/(n^i_S + n^i_M + n^i_L)$, where $n$ is the number of samples. The normalized distribution $R^i_{S,M,L}/\sum_i R^i_{S,M,L}$ shows the cross-shore probability of occurrence per species. In the OBIS data, samples with life stage labels for juveniles or adults only account for a small proportion of all, respectively 20%, 13% and 35% for the surface epipelagic, migrating epipelagic and mesopelagic fish. The cross-shore probability distribution for labeled juveniles or adults fish is thus more variable (Fig. 5D-F) than when all samples are included (Fig. 5A-C). But similarities are visible. Figure 5A,B the cross-shore distribution indicates a decreasing probability of occurrence of small size species as moving offshore, for both the surface epipelagic and migrating epipelagic fish, while larger species tend to distribute more homogeneously along the cross-shore section. For larger species, the probability of occurrence is even slightly higher offshore (Fig. 5A,B). For the mesopelagic fish, most samples include small species (95%), therefore the distribution appears homogeneous regardless the distance to shore, the distribution for medium and large species is then probably spurious (Fig. 5C). We compare this cross-shore succession of the size of occurring species with simulations.

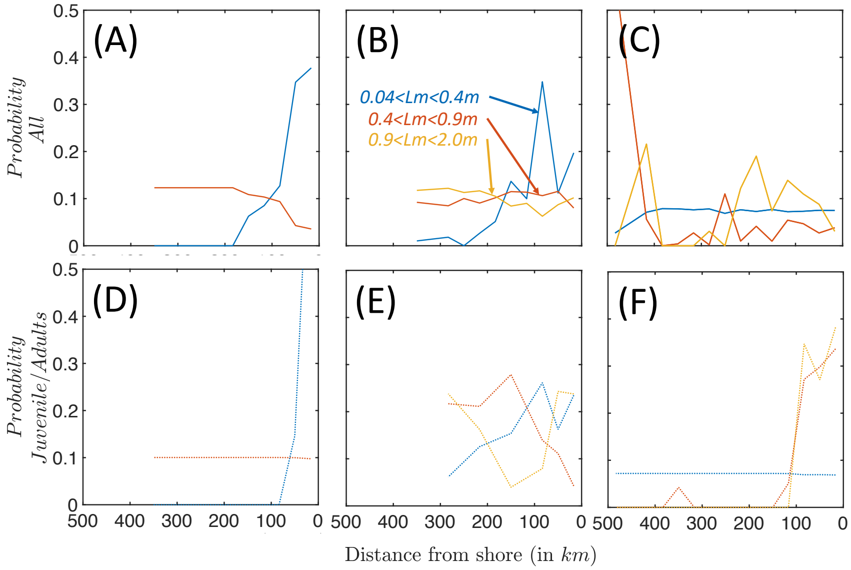


Figure 5: Cross-shelf probability of occurrence per species of increasing size from OBIS: (a-c) when all observation are including, for (A) epipelagic ($n_{obs}^{Epi} = 4088$), (B) migratory epipelagic ($n_{obs}^{Mig} = 10279$) and (C) mesopelagic ($n_{obs}^{Meso} = 11125$) species; (D-F) when only labelled observation of juveniles and adults are including, for (D) epipelagic ($n_{obs}^{Epi} = 803$), (E) migratory epipelagic ($n_{obs}^{Mig} = 1205$) and (F) mesopelagic ($n_{obs}^{Meso} = 3819$) species.

### S9: Eco-regions of the California Current

For tuning of APECOSM-1D and for comparison of the results, we identify 15 eco-regions in our domain of the California Current, based on relevant environmental drivers $l$ provided by the ROMS-BEC forcing. The selected driver are temperature $T$, oxygen $O_2$, biomass of diatoms $B^{Diat}$ and zooplankton $B^{Zoo}$, particle flux at 50m depth $F_{POC}$. With each driver, we generate a map of: (1) the mean value $\overline{l(i,j)}$ (where $i$,$j$ are the coordinates) over years 1997 to 2007; (2) the amplitude of variation $\Delta l = max(l(i,j)) - min(l(i,j))$ from 1997 through 2007; (3) the variability expressed by the standard deviation $std(l(i,j))$. We generate these maps for the epipelagic realm, averaging drivers from surface to 50m depth, for the mesopelagic realm, averaging drivers from 101 to 517m depth.

For the epipelagic and mesopelagic layers, we processed the generated maps through $k$-mean clustering to reveal eco-regions for the epipelagic (Fig. 6A) and mesopelagic (Fig. 6B). We respectively keep 5 and 6 eco-regions the epipelagic and mesopelagic layers. Merging both maps we identify 15 eco-regions accounting for both latitudinal and cross-shore gradients (Fig. 6C)).

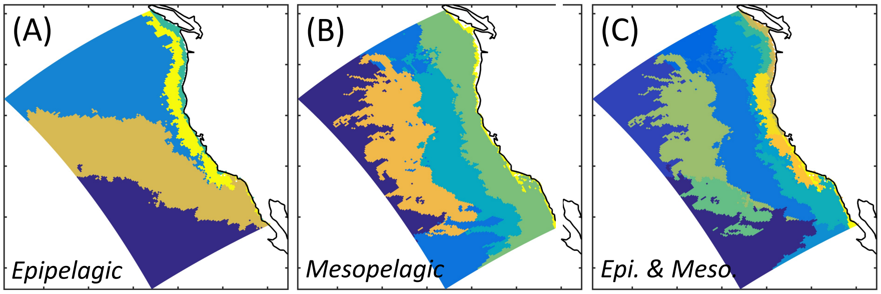


Figure 6: Eco-regions of the California Current: (A) for the epipelagic layer from surface to 50m depth; (B) for the mesopelagic layer from 101 to 517m depth; (C) merging epipelagic and mesopelagic.

### S10: Local biomass budget

The local biomass budget $B$ per grid cell $(i, j)$ is a function of local biomass accumulation through growth $G$, biomass loss through mortality $M$, and in-coming/out-going biomass through the boundaries of the cells $F_{Phys,Swim}$:

$$\frac{dB_{i,j}}{dt} = G_{i,j} - M_{i,j} + \nabla_{i,j} F_{Phys} + \nabla_{i,j} F_{Swim} \tag{13}$$

for each community. We define the metabolic source/sink of biomass from food-web related terms $\Delta_{META} = G_{i,j} - M_{i,j}$. We define the passive source/sink of biomass from the passive biomass transport by currents $\Delta_{CURR} = \nabla F_{Phys}$. We define the active source/sink of biomass from the active biomass transport controlled by swimming organisms $\Delta_{SWIM} = \nabla F_{Swim}$. Each of these indicators is averaged over 8 years of simulation, per grid cell.

### S11: Best APECOSM-1D simulations

The ensemble of APECOSM-1D simulations allows the identification of (1) parameter sets matching observed averaged biomass estimates over the California Current and (2) parameter sets matching regional variations of pelagic and migratory biomass. It also allows studying the sensitivity of predictions to parameters.

Comparing distributions of optimized parameters to the non-optimized distributions using a two-sample Kolmogorov-Smirnov test reveals that $C_{FONC}$, $CRISTAL_{CRIT}$, $M$, and the zooplankton biomass size range control $\overline{B^{Mig}}_{MW}$ and the ratio $\overline{B^{Mig}_{MW}}/\overline{B^{Epi}_{MW}}$ that match observations (see Table 3). Parameters for the size-selective predation $k_{1,2}$ and size ranges for diatoms and particles, show no significant differences from priors.

For the 152 simulations that produce realistic biomass densities, the multi-linear regression between free parameters and the average biomass per community $\overline{B^{Epi,Mig}}_{MW}$, and the ratio $\overline{B^{Mig}_{MW}}/\overline{B^{Epi}_{MW}}$, are used to assess the parameter sensitivity of the model (Table 3). The size-selective predation $k_{1/2}$ influences most of the variability of predicted biomass. This is followed by parameters that influence the community-level mortality, $M$, and the predation intensity, $C_{FONC}$. This predation intensity also appears as the main control of the biomass ratio. Parameters coupling the model with low trophic levels are mostly insignificant, except the average size of the particles that feed the migratory community, $\overline{L^{POC}}$. Prey biomass density is a secondary control, except when prey are scarce in the deep ocean. Otherwise, the schooling parameter $CRISTAL_{CRIT}$ has no significant influence, likely because it influences both mortality ($M$) and predation intensity ($C_{FONC}$). Low $CRISTAL_{CRIT}$ leads to higher mortality as well as more fish production because of enhanced predation, and vice versa for high $CRISTAL_{CRIT}$.

After analysis of the APECOSM-1D simulations, we identify 6 best ensembles whose parameters are summarised Table 4.

Table 3: Sensitivity of APECOSM-1D to the food-web parameters. The Kolmogorov-Smirnov (KS) test shows the similarity between optimised and non-optimised parameter sets. The multi-linear regression between parameters of the 152 optimized parameter sets with $\overline{B_{MW}^{Epi,Mig}}$ show the variation of the simulated biomass for the corresponding parameters, when normalized with zscore. For low trophic levels (i.e. LTL = POC / Diat / Zoo), $\Delta L^{LTL} = log10(L_{max}^{LTL}/L_{min}^{LTL})$, $\overline{L^{LTL}} = log10(\sqrt{L_{max}^{LTL}L_{min}^{LTL}})$. **Bold** font values highlight significantly different parameter distributions, significant predictors ($p < 0.01$).

| | $\log_{10}(C_{FONC})$ | $\log_{10}(CRISTAL_{CRIT})$ | $\log_{10}(M)$ | $k_1$ | $k_2$ | $\Delta L^{POC}$ | $\overline{L^{POC}}$ | $\Delta L^{Diat}$ | $\overline{L^{Diat}}$ | $\Delta L^{Zoo}$ | $\overline{L^{Zoo}}$ |
|---|---|---|---|---|---|---|---|---|---|---|---|
| KS p-value | **5.4 $10^{-5}$** | **4.5 $10^{-37}$** | **1.8 $10^{-72}$** | 0.07 | 0.80 | 0.52 | 0.29 | 0.65 | 0.51 | **5.2 $10^{-3}$** | **5.5 $10^{-3}$** |
| $\log_{10}(\overline{B_{MW}^{Epi}})$ ($R^2 = 0.3$) | **-0.37** | 0.09 | **-0.29** | **-0.39** | **0.45** | 0.03 | 0.08 | -0.03 | 0.06 | -0.15 | 0.10 |
| $\log_{10}(\overline{B_{MW}^{Mig}})$ ($R^2 = 0.37$) | **-0.25** | 0.09 | **-0.30** | **-0.48** | **0.60** | 0.13 | **0.20** | -0.10 | 0.04 | -0.11 | 0.07 |
| $\overline{B_{MW}^{Mig}}/\overline{B_{MW}^{Pel}}$ ($R^2 = 0.17$) | **0.28** | -0.06 | 0.08 | -0.02 | 0.14 | 0.17 | 0.15 | -0.09 | -0.07 | 0.15 | -0.05 |



Table 4: Parameters of the 6 best APECOSM-1D simulations.

| Ensemble | $C_{FONC}$ | $CRISTAL_{CRIT}$ | $M$ | $k_1$ | $k_2$ | $\Delta L_{POC}$ | $\overline{L_{POC}}$ | $\Delta L_{Diat}$ | $\overline{L_{Diat}}$ | $\Delta L_{Zoo}$ | $\overline{L_{Zoo}}$ |
|---|---|---|---|---|---|---|---|---|---|---|---|
| 1076 | 1.03 | 133.2 | 7.8 $10^{-3}$ | 1.16 | 2.05 | 1.89 | $-2.81$ | 1.43 | $-4.57$ | 1.82 | $-2.89$ |
| 1136 | 23.7 | 21.23 | 2.1 $10^{-3}$ | 1.11 | 1.11 | 2.45 | $-3.07$ | 1.14 | $-4.32$ | 1.77 | $-2.93$ |
| 1384 | 30.0 | 0.20 | 1.4 $10^{-3}$ | 1.13 | 1.28 | 1.78 | $-3.07$ | 1.09 | $-4.55$ | 2.20 | $-3.14$ |
| 1721 | 0.93 | 0.4 | 1.4 $10^{-3}$ | 2.44 | 1.14 | 2.22 | $-2.93$ | 1.16 | $-4.69$ | 1.77 | $-2.84$ |
| 1804 | 17.4 | 34.2 | 1.3 $10^{-3}$ | 2.05 | 1.75 | 2.08 | $-2.99$ | 0.90 | $-4.29$ | 1.89 | $-2.92$ |
| 4148 | 1.57 | 1.33 | 1.1 $10^{-3}$ | 2.67 | 1.29 | 1.69 | $-2.86$ | 1.16 | $-4.28$ | 1.96 | $-2.76$ |

### S12: Best APECOSM-CC simulations

Spatial biomass transport influences the biomass density distribution between eco-regions. For 6 sets of parameters that best reproduce biomass density gradients with APECOSM-1D (see S11), we run APECOSM-CC nested into a Pacific wide configuration. Figure 7 shows the total biomass distribution in the California Current ($\overline{B^{Epi+Mig}}$, panels A-F) and the ratio between migratory and pelagic biomass ($\overline{B^{Mig}}/\overline{B^{Epi}}$, panels G-L) for the 6 simulations. For simulations 1136, 1384 and 1804, the ratio between migratory and epipelagic biomass falls out of the expected range. We select simulation 1076 as the best simulation overall as it best reproduces observed variations of biomass density.

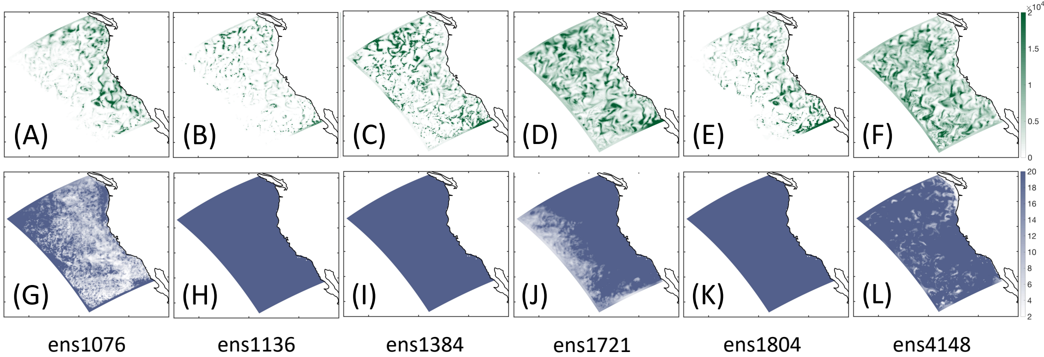


Figure 7: APECOSM-CC simulations for 6 best sets of parameters. (A-F) Biomass density distribution ($\overline{B^{Epi+Mig}}$) in December. (G-L) Ratio between migratory and epipelagic biomass ($\overline{B^{Mig}}/\overline{B^{Epi}}$) averaged over the year.

### S13: Cross-shore primary production

Figure 8 shows the relative distribution of biomass at low trophic levels $B^{LTL}$ from ROMS-BEC when averaged in regular distance bins, averaged over the years 1999 to 2006. While the diatoms and POC dominate the upwelling along the coast, the zooplankton are slightly more homogeneously distributed. Summing all low trophic levels, higher biomass densities are coastal (black line Fig. 8).

When compared with observation of the meso-zooplankton biomass density from MAREDAT averaged over the surface to 200m depth layer (Moriarty and O'Brien (2013)), the simulated zooplankton biomass is more homogeneously distributed than expected (compare plain and doted yellow lines Fig. 8).

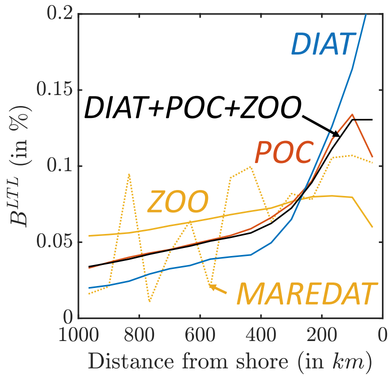


Figure 8: Relative distribution of biomass density at low trophic levels along a cross-shore section, diatoms (in blue), particulate organic carbon (in red), zooplankton (in yellow), the sum of all (in black). The distributions are determined from the average of simulated years 1999 to 2006. The dotted line show the distribution of meso-zooplankton from the MAREDAT database.
